# Integration of eQTL and Machine Learning Methods to Dissect Causal Genes with Pleiotropic effects in Genetic Regulation Networks of Seed Cotton Yield

**DOI:** 10.1101/2023.06.20.545658

**Authors:** Ting Zhao, Hongyu Wu, Xutong Wang, Yongyan Zhao, Luyao Wang, Jiaying Pan, Huan Mei, Jin Han, Siyuan Wang, Kening Lu, Menglin Li, Mengtao Gao, Zeyi Cao, Hailin Zhang, Ke Wan, Jie Li, Tianzhen Zhang, Xueying Guan

## Abstract

Expression quantitative trait loci (eQTL) provide a powerful means of investigating the biological basis of genome-wide association study (GWAS) results and exploring complex traits or phenotypes. In addition to identifying the causal gene in *cis*, eQTL analysis also reveals a large number of trans-regulated genes located on different chromosomes, which form a gene regulatory network (GRN) that complements the GWAS locus. However, the dissection of a GRN and the crosstalk underlying multiple agronomical traits, along with prioritizing important genes in eQTL-derived GRNs, remains a major challenge. In this study, we generated 558 transcriptional profiles of lint-bearing ovules at one day post-anthesis (DPA) from a selective core cotton germplasm, from which we identified 12,207 eQTLs. By integrating with a GWAS catalog, we found that 66 out of 187 (35.29%) known phenotypic GWAS loci are colocalized with 1,090 eQTLs, forming 38 major functional GRNs predominantly (30 out of 38) associated with seed size-related phenotypes. Of the eGenes, 34 were shared between at least two functional GRNs, exhibiting pleiotropic effects, such as *NF-YB3*, *GRDP1*, and *IDD7*. Narrow-sense heritability analysis showed that the heritability increased with combining the eQTLs with GRNs compared to those with previous yield trait GWAS loci. The extreme gradient boosting (XGBoost) machine learning approach was then applied to predict seed cotton yield phenotypes based on gene expression. Top-ranking eGenes (*NF-YB3*, *FLA2*, and *GRDP1*) derived by XGBoost with pleiotropic effects on yield traits were validated, along with their potential roles by correlation analysis, domestication selection analysis, and transgenic plants. This study provides insights into the mining of GRNs in relation to the pleiotropy of phenotype. The combination of eQTL and machine learning approaches is efficient in improving the genetic dissection of agricultural traits.

## Introduction

The genome wide association study (GWAS) is a common method for detecting associations between genetic variation and phenotype. The GWAS can be traced back to the first decade of the 2000s, ^1, 2^ or earlier. To date, tens of thousands of associated loci have been cataloged in major crops such as rice ^3^, wheat ^4^, maize ^5^, and cotton ^6^. However, while GWAS has been very successful in identifying loci associated with phenotypes, it still suffers from major limitations when it comes to pinning down causal genes due to issues with population structure, the missing heritability of rare variations, effects from *trans* regulation networks, and more besides.

Notably, although accumulated common genomic variants with small effect sizes can contribute to various traits, they may be filtered out from GWAS results by a stringent significance threshold ^7, 8^. In addition, GWASs focus on genomic variations in the form of common SNPs and neglect rare variations with minor allele frequency (MAF) values of less than 1-5% ^9^. More importantly, GWASs are limited in their ability to pinpoint causal variants and candidate genes due to the resolution of linkage disequilibrium (LD) in small population sample size. LD blocks can range in size from 30 kb in a common maize population ^10^ to ∼100 -200 kb in cultivated rice ^3^ and ∼300-500 kb in cotton ^11, 12^. Consequently, the candidate genes associated within a LD can range in number from a few to a few hundred. Theoretically, any gene within the LD block of a GWAS locus could not be excluded as having a causal effect on the corresponding trait. Additionally, the majority of GWAS loci are located in non-coding region and likely manifest their effects by regulating distant gene expression in *trans*^13^. Unfortunately, GWAS cannot accommodate the direct identification of genome-spanning gene regulation networks (GRNs). Collectively, these limitations prevent further application of GWAS in navigating the critical step of hub gene selection for precision genome editing in the interest of crop improvement.

One solution for overcoming the limitations of GWAS is to integrate relevant expression data in addition to genetic variations. Notably, gene expression changes are efficient at introducing phenotypic changes in crops ^14^. Analysis of expression quantitative trait loci (eQTL) is a method of establishing connections between genetic variants and gene expression by identifying expression-associated SNPs (eSNPs) and their associated genes (eGenes). eGenes can be located either within the same region as an eSNP or in a distal region, there eQTL analysis can detect associated eGenes in *cis* and also in *trans*. Alternatively, multiple genes can be regulated by a single *trans-* eSNP, termed as eQTL hot-spot, module or GRN ^15^. Therefore, each GRN is composed of eGenes both in *cis* and *trans*. The latest study based on the Genotype-Tissue Expression (GTEx) dataset (v.8) demonstrated that a median of 21% of GWAS loci from 87 tested complex traits colocalized with a *cis*-eQTL when aggregated across 49 tissue types ^16^. Gene-gene interactions within a GRN are proposed to be components of the missing heritability for complex traits ^17^. The size of GRNs, termed as eQTL hotspots, ranges widely. For example, cotton contains 243-1,300 genes in its GRNs, while maize has 125. ^18–20^. Thus, although integrated eQTL and GWAS analysis can provide functional GRNs associated with phenotypes, prioritizing impactive genes in the GRN and their power to affect the phenotype is still a challenge.

Extreme gradient boosting (XGBoost) is a machine learning method, specifically a type of decision tree ensemble model for classification and regression modeling (Chen, 2016). It is remarkable for its ability to process missing data efficiently and flexibly and to assemble weak prediction models from which can be built an accurate one ^21, 22^. In a competition hosted by Kaggle.com, XGBoost was found to be the best algorithm for machine learning and prediction ^23^. As it can evaluate the degree of feature importance, XGBoost can be used to prioritize genes according to their criticality, as has been reported in human populations ^24, 25^. Pioneering research has been conducted in plants, specifically mining N-responsive genes in *Arabidopsis thaliana* and maize ^26^, however, the application of XGBoost to populations of crops or other plants is still in a preliminary stage.

Previous GWASs in cotton have revealed associated loci for multiple important agronomic traits including fiber production ^6, 12^, seed cotton yield ^27, 28^, fiber quality ^11, 29^, and abiotic stress tolerance ^30–32^. Other recent studies have employed eQTL hotspot analysis methods to dissect the genetic GRNs that regulate fiber quality traits and pollen sterility ^19, 33^. Nonetheless, while a large number of phenotype-associated loci have been reported in cotton populations, the causal genes and other important genes within functional GRNs are still largely unknown due the lack of data mining methodology.

To characterize the genetic basis of cotton seed size and yield, identify causal genes, and elucidate the underlying GRNs associated with GWAS loci, we designed an integrative eQTL and GWAS analysis using transcriptomes from the <u>C</u>hina <u>u</u>pland <u>c</u>otton <u>p</u>opulation, CUCP1. The identified eGenes clustered into 38 functional GRNs based on the colocalization of expression-associated lead SNPs (eSNPs) and phenotype-associated lead SNPs (pSNP) within LD blocks. The joint additive effect of yield-related GRNs was validated based on narrow-heritability. Using an XGBoost-derived featurex’x importance ranking, the causal genes *NF-YB3, FLA2* and *GRDP1* from GRN were validated as having functional impacts on seed development.

## Results

### Study overview

**Figure. 1** illustrates the aim of this work, which is to construct gene regulation networks (GRNs) and mine the genes that are important for seed size and fiber yield.

**Figure 1:**
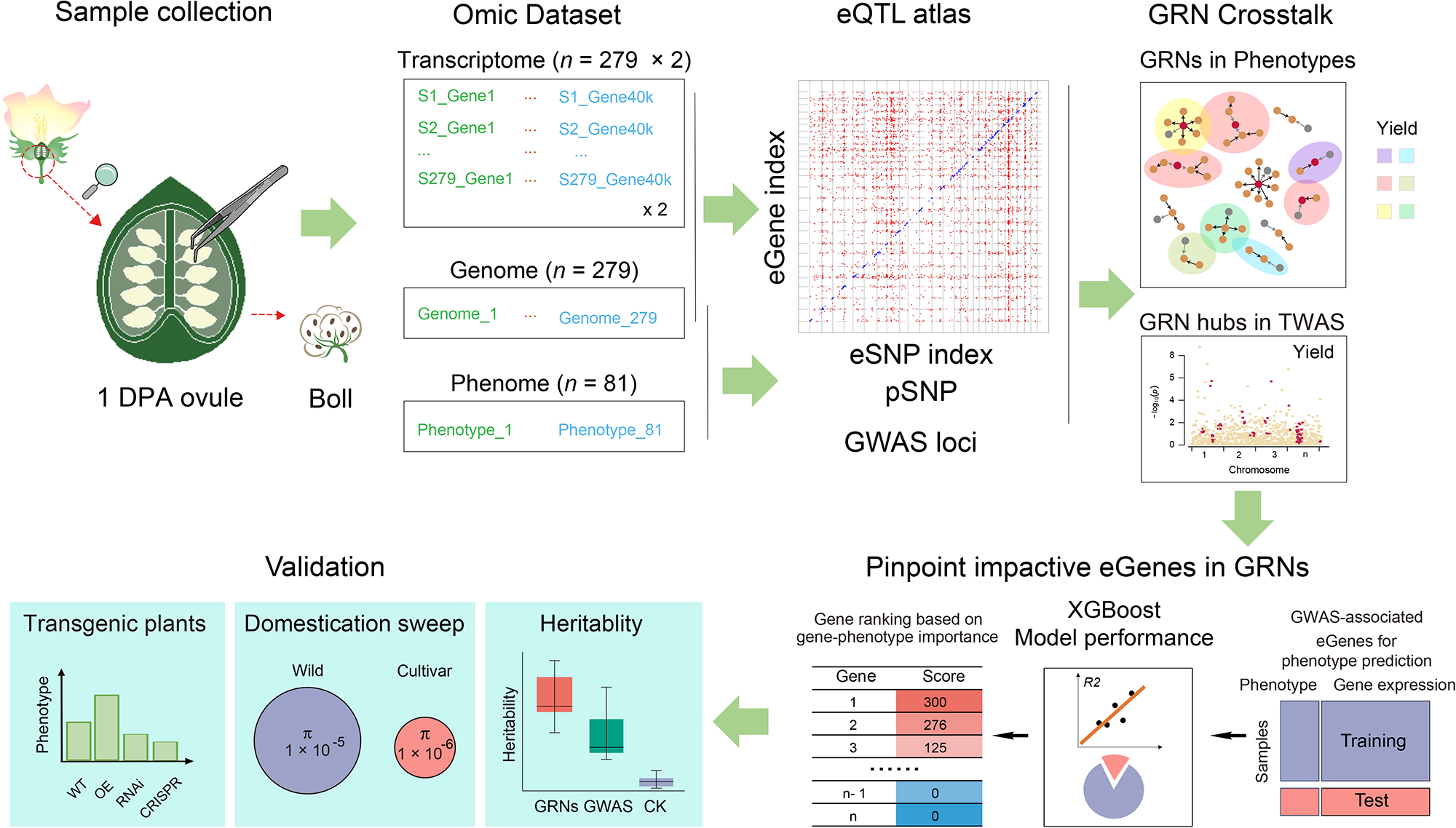
Graphic summary of datasets and analyses performed in the present study. The principal goal is to dissect the genetic networks underlying phenotypic correlations and mine impactful genes. This schematic chart represents our datasets and methods for network dissection and prioritization of impactful eGenes by integrating multiple omics (transcriptome, genome, and phenome).

#### Data

A core germplasm was collected for the <u>C</u>hina <u>u</u>pland <u>c</u>otton <u>p</u>opulation, named CUCP1, comprising a total of 279 *Gossypium hirsutum* accessions, including 34 wild/landrace accessions and 245 cultivated accessions. The collection of cultivated accessions is adapted from our previous GWAS catalog (**Table S1**) ^12^.

Cotton fiber differentiates from ovule epidermis at about -1 to 1 day post-anthesis (DPA). Approximately 25-30% of the epidermal cell can differentiate into fiber cells, which largely determine the fiber yield on each seed ^34, 35^. To understand the expression variation associated with genetic variation in CUCP1 at the fiber-yield determining stage, we profiled the transcriptomes of 1-DPA lint-bearing ovules for all 279 accessions, with two biological replicates (**Figure 1**). The transcriptomes collectively provided 13.82 billion (mean = 24.90 million per sample) paired-end reads, with an average 97.11% rate of unique mapping (cultivar 97.23%; wild 96.32%) to the TM-1 cotton reference genome ^36^ (**Table S1**). According to previous reports, most GWAS loci map to non-coding regions and potentially point to non-coding variants ^13^. Accordingly, to detect as many causal genes as possible, we also annotated long non-coding RNAs (lncRNAs) in non-coding regions and quantified their transcription for eQTL mapping (***Materials and Methods***). A total of 37,108 protein-coding genes (PCGs) and 6,251 lncRNAs met the criteria for expressed genes (***Materials and Methods***), accounting for 50.99% of all annotated genes in the TM-1 upland cotton reference genome ^36^; These were used for further analysis. In parallel, whole-genome sequencing (WGS) of the accessions generated a total of 1,186,673 biallelic high-quality SNPs (minor allele frequency > 0.05 and missing ratio < 20 %), which were used for eQTL mapping (**Figure 1**) ^12^.

#### Workflow

(1) GWAS and eQTL were integrated to obtain phenotype-associated lead SNPs (pSNPs) and gene expression-associated lead SNPs (eSNPs), respectively. SNPs within the same LD block (*r*^2^ > 0.1) were clustered. Accordingly, their associated eGenes were grouped as GRNs. GRNs with colocalized pSNP were considered as functional GRNs. (2) The eGenes in functional GRNs was used as the feature for the XGBoost algorithm in phenotype regression prediction. The model’s performance was evaluated using Pearson correlation coefficient (*PCC*). Next, the eGenes were ranked according to the feature importance score exported from the model. (3) The top-ranked eGenes were selected for functional validation using heritability analysis, domestication sweep identification, and transgenic plants (**Figure 1**).

### A map of eQTLs associated with fiber-bearing ovule development

The quality of the 558 (279 × 2) transcriptomes and their applicability to eQTL mapping were evaluated by calculating Pearson correlation coefficients (*PCCs*) from the transcriptome profiles. The PCCs of the two biological replicates (mean *r* = 0.93) were found to be significantly higher than those of different accessions (mean *r* = 0.77, *P* < 1 × 10^-16^, Mann-Whitney test) (**Figure 2A**). Principal component analysis (PCA) was also used to reveal the genetic similarities and differences of expression patterns (**Figure 2B**).

**Figure 2:**
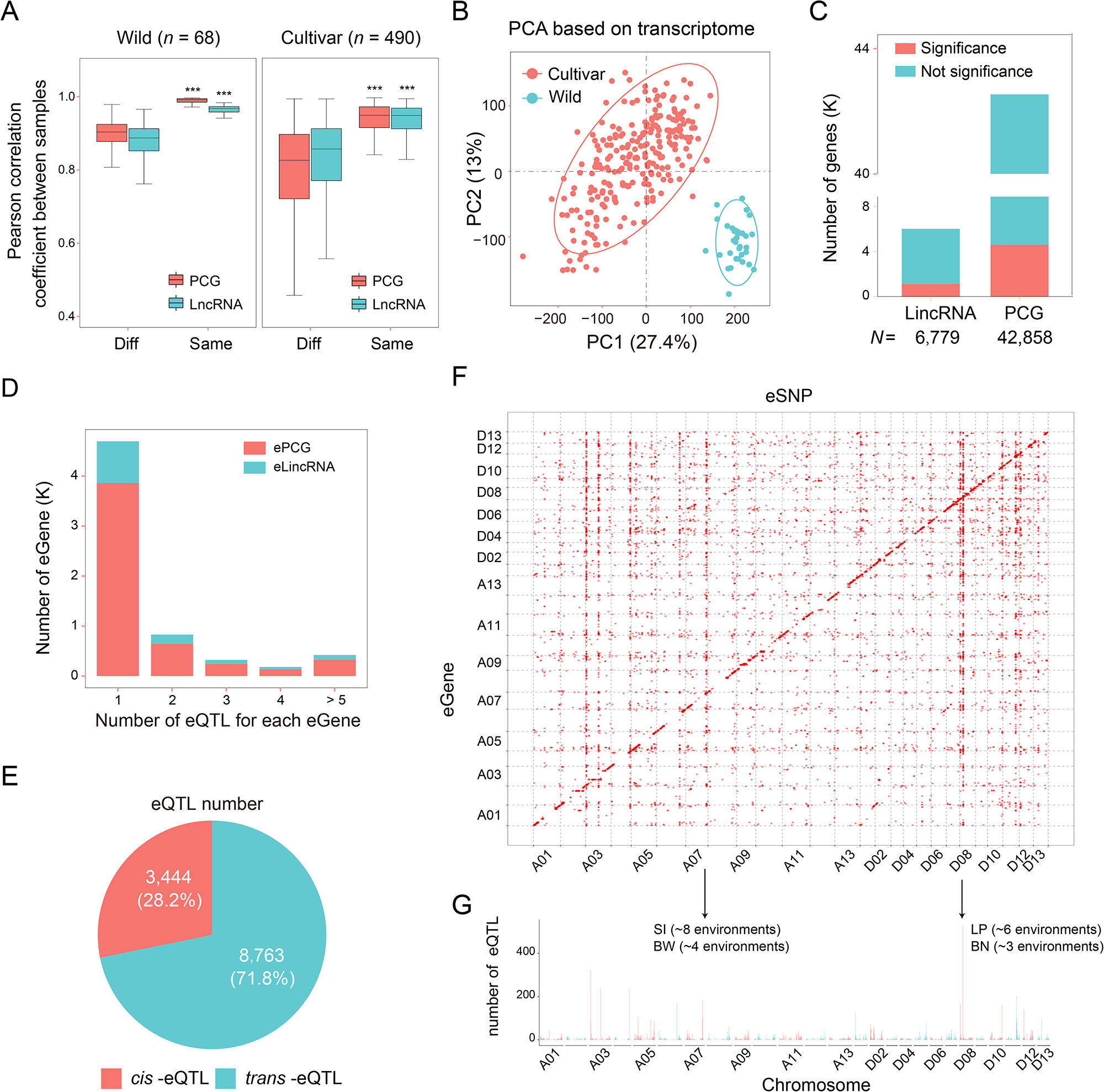
eQTL map for one-day post anthesis (DPA) lint-bearing ovules from 588 samples. (A) Pearson correlation coefficient (PCC) of samples based on protein coding gene (PCG) and lncRNA expression quantifications; correlations compare replicates of the same accession (Same) and randomly selected samples from different accessions (Diff). Quantifications were normalized to FPKM before calculating the pair-wise PCC. Box plot shows the median and interquartile ranges (IQR). The end of the top line is the maximum or the third quartile (Q) + 1.5× IQR. The end of the bottom line denotes either the minimum or the first Q – 1.5× IQR. Dots are either more than the third Q + 1.5× IQR or less than the first Q – 1.5× IQR. (*** *P* < 0.001, two-tailed Mann-Whitney test). (B) Distinct separation of wild and cultivar groups was observed with principal component analysis (PCA) of transcriptome profiles. (C) Numbers of lncRNAs and PCGs associated with eQTLs. (D) Number of eQTLs mapped for each eGene. The x-axis represents the number of eQTLs mapped for each eGene, and the y-axis represents the number of eGenes in each group (PCG and lncRNA). (E) Pie chart showing the number and proportion of *cis*- and *trans*-eQTLs. (F) Scatter plot of 8,088 high-confidence eSNP-expression associations, with expression of 6,449 eGenes (y-axis) against 12,207 eQTLs. Each dot represents a detected eQTL. (G) eQTL hotspot distribution across the genome, was determined in 1-Mb windows. The y-axis indicates number of eGenes and is plotted against genetic location. Arrows indicate two eQTL hotspots co-located with cotton yield GWAS loci. SI, seed index; BW, boll weight; BN, boll number; LP, lint percentage.

eQTL mapping was subsequently performed with Efficient Mixed Model Analysis Expedited (EMMAX) ^37^ using the obtained SNPs and expression profiles. A total of 12,207 eQTLs were detected, involving 8,088 eSNPs and 6,449 eGenes (PCG *n* = 5,197, lncRNA *n* = 1,252), under a suggested threshold of *P* < 2.18 × 10^-6^ (**Figure 2C; Table S2**)^19^. An average of 1-2 eQTLs were mapped for each eGene (**Figure 2D**), suggesting that the expression variation is under relatively simple genetic control. The mapped eQTLs were further classified as *cis* or *trans* according to relative eGene location using an empirical value^38, 39^ (that is, the SNP was within ±1 Megabase [Mb] of the transcription start or termination sites [TSS or TTS] of each gene), yielding 3,444 *cis* eQTLs (involving 1,185 eGenes) and 8,763 *trans* eQTLs (involving 5,869 eGenes) (**Figure 2E**). For *cis*-eQTLs, the associated lead eSNPs were predominantly distributed in adjacent genes and enriched in proximity to TSSs or TTSs (**Figure S1**). *Cis*-eQTLs showed higher association than *trans*-eQTLs did (Mann-Whitney test, *P* < 2.2 × 10^-16^) (**Figure S2A**). More than 77% of the eQTLs were shared between the two biological replicates (**Figure S2B; Table S2**). In addition, 21.42% of eQTLs showed the variance between the wild and cultivated accessions (**Figure S2C-D**). The distribution patterns and frequencies of eQTLs reported here are consistent with previous reports in maize seedlings ^20^, *Brassica napa* seeds ^40^, rice shoots ^41^, cotton 15 DPA fibers ^19^ and *Arabidopsis* shoots ^42^. With regard to chromosomal location, the eQTLs identified here exhibited a significantly disproportionate distribution, forming 293 eQTL hotspots (**Figure 2F; Table S3**)The most notable hotspots spanned 1,756 kb (from 88,974,035 bp to 90,730,903 bp) on chromosome ChrA07 and 7.23 kb (from 2,899,413 bp to 2,906,643 bp) on D08.They were determined to overlap with stable GWAS loci (**Figure 2G; Table S3**).

### Phenotypic Relevance of Gene Regulation Networks Derived from eQTL

To systematically characterize the GRNs derived from eQTL analysis, eGenes either in *cis* or *trans* associated the eSNPs within the same LD block (*r*^2^ > 0.1) were grouped as one GRN. This yielded 1,014 GRNs (**Figure 3A**), with the number of eGenes in each GRN ranging from 2 to 527 with the average of 13 (**Figure S3**).

**Figure 3:**
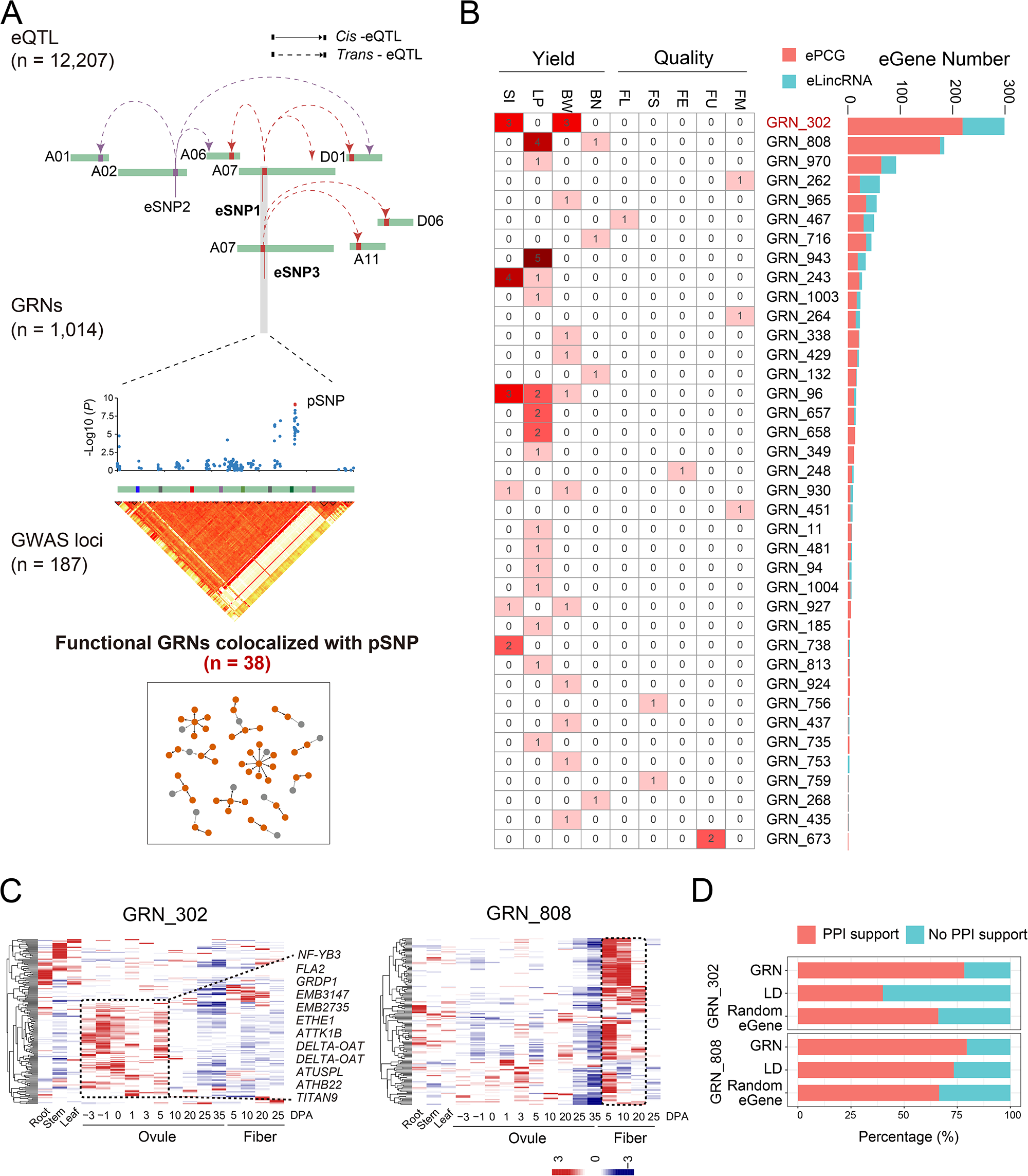
Gene regulation networks (GRNs) with phenotype-associated feature SNPs. (A) Analytical workflow for functional GRN construction. Both GWAS and eQTL analysis were conducted to obtain phenotype-associated lead SNPs (pSNPs) and gene-expression associated lead SNPs (eSNPs), respectively. Those eSNPs/pSNPs within the same LD block (*r*^2^ > 0.1) were merged into one lead SNP, and eGenes within an LD block were clustered into a GRN. GRNs having feature pSNPs were considered to be functional GRNs. (B) Heatmap showing the 38 functional GRNs and their genotypic associations. Numbers in boxes indicate how many times the lead causal SNP satisfied the *P-*value threshold for genome-wide significance in the GWAS. The bar plot in the right panel shows the number of PCGs (red) and lncRNAs (blue) in each GRN. (C) Heatmap showing expression of eGenes in GRN_302 and GRN_808 across different tissues. Thirteen eGenes reported to play roles in seed and embryo development are highlighted. (D) Bar plot showing the percentage of eGenes in GRN_302 and GRN_808 that are also identifiable as interactors in the STRING protein-protein interaction database (https://cn.string-db.org/). Randomly selected eGenes and genes in LD blocks were used as controls.

The set of phenotype-associated SNPs (pSNPs) were adapted from the previous GWAS catalog and represents 187 GWAS loci ^12^. The best linear unbiased prediction (BLUP) values for each of trait were also calculated based on phenotypic data from nine environments (**Table S4**). These values segregated the genetic and environmental effects that influence the phenotypes. The association signals identified by BLUP and SI trait were consistent with those GWAS loci identified by the phenotype in 2007, 2008, and 2009, respectively (**Figure S4**). With the above analysis, the pSNP used in this study were reliable.

To determine whether the obtained GRNs tended to have functional consequences associated with phenotype, the pSNP and eSNP on the same LD were defined as being colocalized with each other. Accordingly, 38 GRNs (3.75%, 38 out of 1,014) were found to colocalize with GWAS loci with the associated function ((**Figure 3A**, **Table S5**); In total, these involved 1,090 eQTL and 701 non-redundant eGenes (**Figure 3B; Table S5**).

Among these 38 functional GRNs, 30 GRNs containing 657 eGenes were related to yield phenotypes, while the other 8 GRNs containing 44 eGenes were related to fiber quality phenotypes (**Figure 3B; Table S5**). Particularly, it was notable that GRN_302 and GRN_808 were associated with yield phenotypes, which together dominant (60.36%) the total eGenes in the functional GRNs (**Tables S5 and S6**). In detail, GRN_302 contained 230 eGenes and its feature eSNP on chromosome A07 (A07:90680544) was co-localized with a GWAS locus represented by a pSNP associated with SI (seed index, weight of 100 seeds) and BW (boll weight) (**Figure 3B; Table S5**), while GRN_808 contained 169 eGenes and its the feature eSNP on chromosome ChrD08 (D08:2903486) was co-localized with a GWAS locus associated with LP (lint percentage) and BN (boll number) (**Figure 3B; Table S5**).

Previous studies have proposed that eGenes within the same GRN should exhibit similar expression patterns in unique cell types relevant to the phenotype of interest ^43^. To confirm whether the eGenes in the same GRNs identified here share similar biological functions, their transcriptional activity was examined using previously published transcriptome profiles from 17 tissues in the upland cotton accession TM-1, which encompassed all developmental stages of seed and fiber ^44^. For both GRN_302 and GRN_808, the eGenes exhibited a general trend of tissue-specific expression (**Figure 3C**). Specifically, over 60% of eGenes in GRN_302 were highly expressed in early ovule/seed development (around 0 to 5 DPA) (**Figure 3C**), while most eGenes in GRN_808 showed similar expression pattern in early ovule and fibers (**Figure 3C**). In addition, 13 eGenes within GRN_302 were found to be homologous to genes with reported roles in seed and embryo development, such as *FLA2* ^45^, *EMB3147*, and *EMB2735* (**Figure 3C**). Within GRN gene-gene interactions were further examined using a protein-protein interaction (PPI) network database ^46^. About 78.33% of the genes in GRN_302 and 79.50% of those in GRN_808 were supported as interactors by this PPI data. These proportions are higher than among randomly selected eGenes and genes in same LD (**Figure 3D**). The above analysis confirmed the eGenes in the GRNs revealed here are highly likely associated with seed development, with potential protein-protein interactions.

### Crosstalk between functional GRNs involved in seed cotton yield

In the present study, each eGene was mapped with 1-2 eQTLs (**Figure 2D**), suggesting that eGenes can be associated with more than one genetic variation. Thus, pairwise comparisons were performed to identify eGenes shared by multiple functional GRNs. This yielded 18 instances of shared eGenes, indicative of crosstalk between different functional GRNs (**Figure 4A**). For example, the dominant network GRN_302 shared eGenes with GRN_96 and GRN_243 (**Figure 4A**); all three of these GRNs were associated with yield phenotypes at significance levels below the threshold of *P* < 2.18 × 10^-6^. Specifically, GRN_302 was associated with SI and BW, while GRN_96 and GRN_243 were associated with SI and LP (**Figure 4B**).

**Figure 4:**
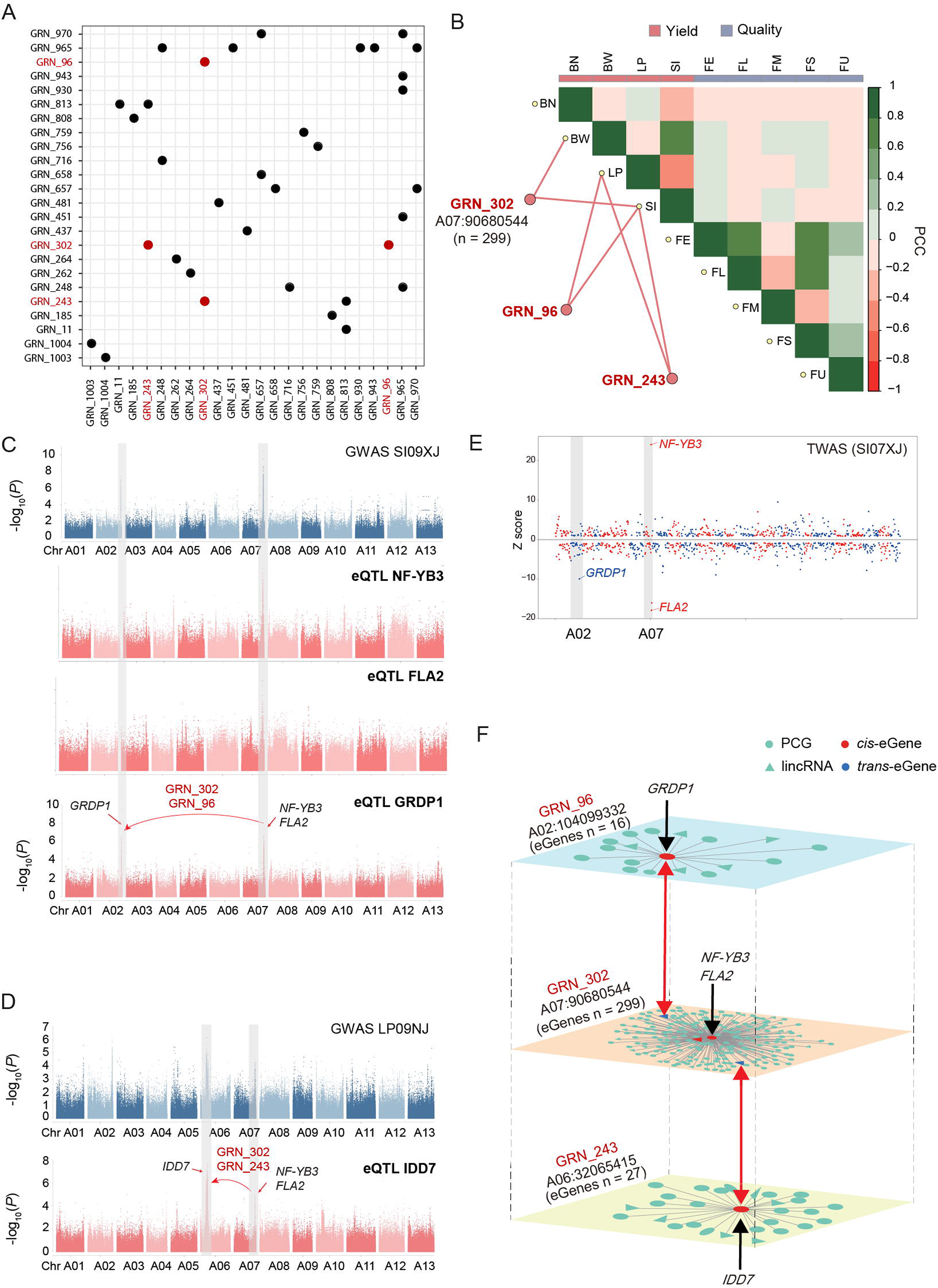
The joint additive effect of yield-related GRNs. (A) Overlap of eGenes across different functional GRNs. The GRNs that shared genes with GRN_302 are colored red. (B) The relationship between representative GRNs (GRN_302, GRN_96, and GRN_243) and their associated phenotypes. The heatmap shows the Pearson correlation coefficients (PCC) of different phenotypes. (C) Manhattan plot for various traits and gene expression; from top to bottom, SI phenotype and expression of *FLA2*, *NF-YB3*, and *GRDP1*. The x-axis is the SNP chromosomal location of SNP and the y-axis is the strength of the association (-log10 (*P*-value)). The causal variations of GRN_302 and GRN_96 were highlighted. (D) Manhattan plot for various traits and gene expression; from top to bottom, LP phenotype and expression of *IDD7.* The x-axis is the SNP chromosomal location and the y-axis is the strength of the association (-log10 (*P*-value)). The causal variations of GRN_302 and GRN_243 are highlighted. (E) Manhattan plot of TWAS results for the SI phenotype. Each point represents a single *cis-*eGene. Genes whose expression is positively or negatively correlated with SI are plotted above or below the black bold line. The genomic positions of each eGene are plotted on the x-axis. *NF-YB3, FLA2, GRDP1*, and *IDD7* were highlighted. (F) Connections across GRN_302, GRN_243, and GRN96. Nodes represent eGenes (circles, PCGs; triangles, lncRNAs). The *cis*-eGenes in each GRN are highlighted in red. Lines represent *trans* regulation.

Regarding specific genes in the regulatory networks, *NF-YB3 (GH_A07G2187)* and *FLA (GH_A07G2189)* are two *cis*-eGenes in GRN_302 (**Figure 4C and Figure S5**). A Manhattan plot showed that the genomic variants within the corresponding GWAS locus were significantly associated with the expression of *NF-YB3* and *FLA2* in all examined environments (**Figure 4C**; **Figures S6 and S7**). *GRDP1* is a *trans*-eGene in GRN_302, but a *cis*-eGene in GRN_96 (**Figure 4C**). eQTL mapping of *GRDP1* (*GH_A02G1719*) revealed two regulatory variants located in two SI-GWAS loci on ChrA02 and ChrA07, which represent *cis-* and *trans-* regulation patterns in GRN_96 and GRN_302, respectively. This finding indicates an interaction between GRN_302 and GRN_96 in controlling SI (**Figure 4C**). Similarly, *IDD7* (*GH_A06G0949*) is a *trans*-eGene in GRN_302, and a *cis* -eGene in GRN_243 (**Figure 4B**). eQTL mapping of the *cis-*eGene *IDD7* likewise identified two regulatory variants located in two LP-GWAS loci in ChrA06 and ChrA07, which represent *cis-* and *trans-* regulation patterns in GRN_243 and GRN_302, respectively (**Figure 4D**).

Here we also adopted the transcriptome-wide and regulome-wide association studies (TWAS) ^47^ to validate the presumed causal role of the *NF-YB3, FLA2*, and *GRDP1*. In total, 297 expression-phenotype associations were found, involved with 83 PCGs, and 13 lncRNAs (*P* < 9.2 × 10^-4^) (**Figure 4E; Table S7**). Consistent with the integrative functional GRNs*, NF-YB3, FLA2*, and *GRDP1* all achieved significance in the TWAS on seed cotton yield traits (**Figure 4E; Table S7**).

Together with the shared eGenes from different GRNs, whether *cis*-or *trans-*, these GRNs might form an interactive connection that collectively constitutes a potential network regulating seed cotton yield (**Figure 4F**).

### Prioritizing the impactful genes in GRNs using XGBoost

The power of each eGene in phenotype regulation is still unclear. Here we employed the XGBoost algorithm to prioritize the most impactful eGenes. Taking the screened eGenes within major functional GRNs as features, XGBoost was used to construct regression models for the phenotype in each environment (**Figure 5A**). The mean Pearson correlation coefficient (PCC) between the true values from the test data and the predicted values was high in SI and LP (**Figure 5B**; **Table S8**). FL (fiber length) and FS (fiber strength) phenotypes were determined as controls, for which the obtained *r* values were lower than 0.1 (**Figure 5B; Table S8**). This confirms the impacts of the identified GRNs on seed cotton yield.

**Figure 5:**
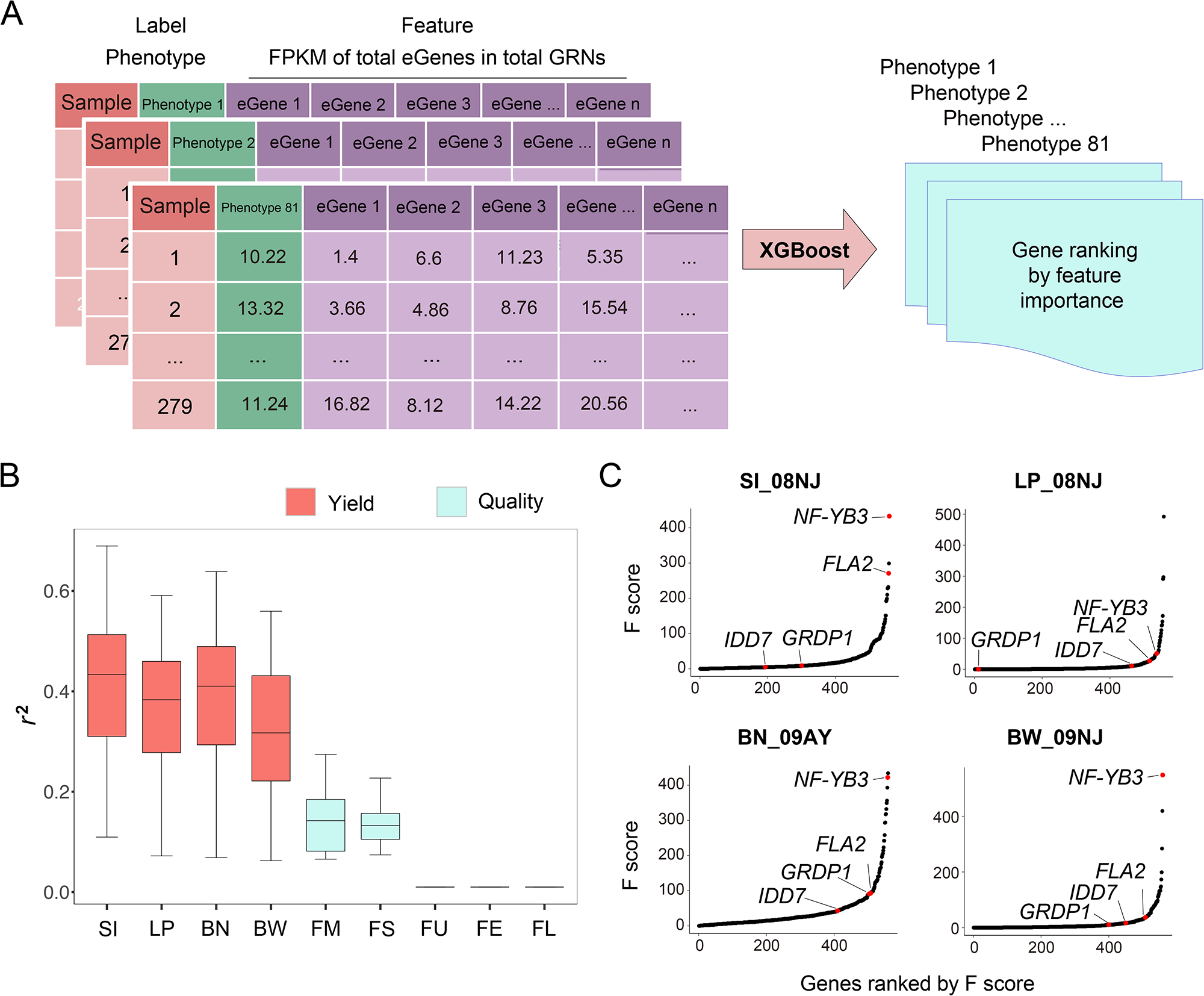
Prioritizing impactive genes in GRNs using XGBoost. (A) Machine learning workflow. The input data consisted of instances (samples) with labels (phenotypes) and values of features (eGenes). Instances were first split into training and testing sets. The training set was further split into a training subset (90%) and validation subset (10%) in a five-fold cross-validation scheme. After tuning the model parameters, the optimal model was used to provide performance metrics based on PCC *r* between the predicted and actual values in each environment, predict labels in the testing set for model evaluation purposes, and obtain feature importance scores. (B) Box plot showing performance based on PCC *r* between the predicted and actual values in each phenotype. (C) Coefficients are averaged from 100 iterations of model building. (D) Feature importance (F score) of each eGene. The x-axis represents genes ordered by F score within each phenotype and the y-axis the F-score value exported by XGBoost.

Next, the feature importance for measuring the distinction in prediction was exported from the XGBoost model (**Table S9**). Among the impactful genes so identified, the potential pleiotropic genes *NF-YB3*, *FLA2*, *GRDP1*, and *IDD7* ranked at the top (**Figure 5D**).

### The interactive GRNs captured the missing heritability for seed size

Together with the shared eGenes, the GRNs form an interactive connection that comprises a potential regulatory network for seed cotton yield. In validating the effects of this extended network, a key question to answer is whether the detected GRNs can provide quantitative power to increase the heritability. To quantify the relative genetic contribution of each GRN to phenotypic variation, narrow-sense heritability (*h*^2^) can be calculated using the local variants ^48^. Hereafter, *h*^2^_GRN_ indicates the local variance explained by SNPs corresponding to *cis*-and *trans-*eGenes within a GRN. In the interests of comparison, we also determined the variance explained by GWAS loci and randomly selected eSNPs, denoted as *h*^2^_GWAS + random_, and that explained solely by randomly selected eSNPs, termed *h*^2^_random_.

The combined heritability of the effect of GRN_302 on SI was found to be significantly increased by about three-fold (19.34%) compared with that of GWAS loci alone (7.00 %), while the heritability of the same number of randomly selected eGenes is 1.84% (**Figure 6A, Figure S8**). The higher heritability of GRN_302 is probably due to the two *trans*-eGenes associated with the causal variants from GRN_96 and GRN_243 (**Figure 4F**). To test this hypothesis, the combined heritability estimated for GRN_302+GRN_96+GRN_243 were evaluated, the result showed that the combined heritability achieved an even higher level of significance when compared with GWAS loci comprised of the causal variants of GRN_302, GRN_96, and GRN_243 (*h*^2^_GRN302+GRN_96+GRN243_, 20.81%; *h*^2^_GWAS_, 8.86%; *h*^2^_random_, 1.50%) (**Figure 6B; Figure S9**), indicating that GRNs can capture phenotype-associated genes that have minor effects and are undetectable by GWAS. For the seed size associated phenotypes of SI, LP, BW, and BN, *h*^2^_GRN_ was significantly higher than either *h*^2^_GWAS, random_ or *h*^2^_random_ (Mann-Whitney *P* = 2.2 × 10^-16^) (**Figure 6C; Figure S9**). Meanwhile, the joint effects of GRNs as represented by *h*^2^_GRN_ did not affect fiber quality traits (FS, FL, FU, and FM), which is consistent with the GWAS results (**Figure 6C**; **Figures S9 and S10**). This control demonstrated the GRNs reveal at 1-DPA ovule stage employing eQTL network predominantly explain the regulation on seed development and seed size.

**Figure 6:**
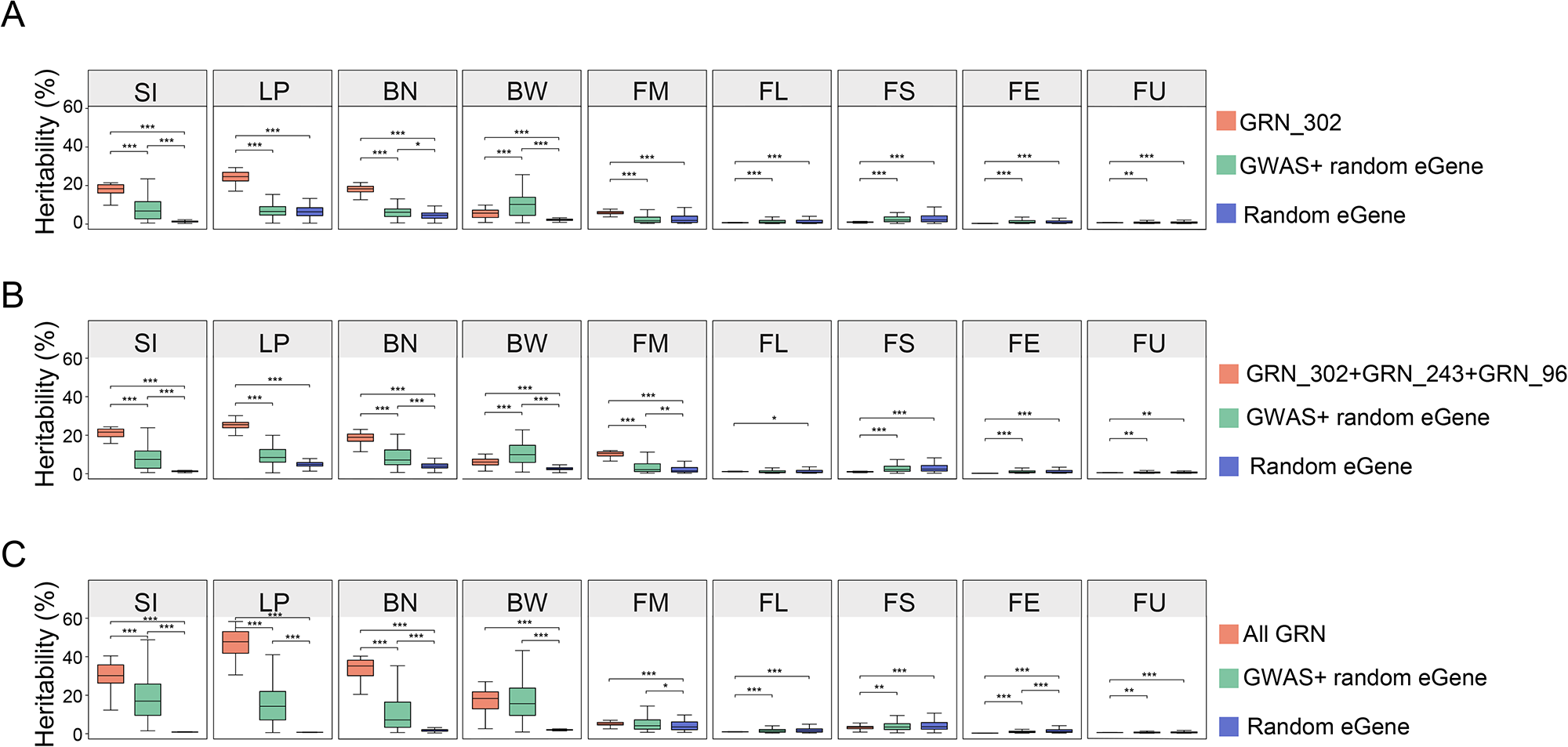
GRNs explained a larger fraction of heritability than the functional GWAS loci. (A-C) Box plots show the estimated heritability (*h*^2^) across different phenotypes explained by different sets of eSNPs shown in the corresponding color code.Boxes show the medians and IQRs. The end of the top line is the maximum or the third quartile (Q) + 1.5× IQR. The end of the bottom line denotes either the minimum or the first Q − 1.5× IQR. Dots indicate values outside those bounds. (*** *P* < 0.001, two-tailed Mann-Whitney test).

### *NF-YB3*, *FLA2*, and *GRDP1* are highlighted as the genes most plausibly governing the seed size phenotype

*NF-YB3, FLA2, and GRDP1* were top-ranked eGenes prioritized by machine learning and TWAS (**Figure 5C**). The feature eSNP on GRN_302 is located within a GWAS locus for which the LD block spans a 270.74-kb region on chromosome A07 and contains 11 annotated genes (**Figure S5**). *NF-YB3* and *FLA2* are two *cis-*eGenes on this locus (**Figure 4C and Figure 7A**). The expression of *NF-YB3* and *FLA2* was significantly correlated with SI and BW phenotypes (**Figure 7B**; **Figures S11 and S12**). The homolog of *NF-YB3* in *Arabidopsis* is *AT4G14540,* which encodes a nuclear factor Y transcription factor, NF-YB3, that has been well-studied in embryogenesis and seed development ^49–56^. The ectopic expression of cotton *NF-YB3* decreased the seed size in the transgenic *Arabidopsis* (**Figure S13**), which confirmed its direct impact on seed development. In addition, these data confirmed the eQTL analysis can efficiently navigate to seed regulation genes. The homolog of *FLA2* in *Arabidopsis* is *AT4G12730*, which encodes fasciclin-like arabinogalactan-protein 2 with reported function in seed development ^45, 57^. *FLA2* is actively expressed in the leaf, flower, and early ovules during seed development in both *Arabidopsis* and cotton (**Figure S14**). Statistical analysis further revealed that the expression of these two eGenes to be negatively correlated, suggesting that *NF-YB3* and *FLA2* may play antagonistic roles in coordinating seed development (**Figure 7C**; **Figures S11 and S12**).

**Figure 7:**
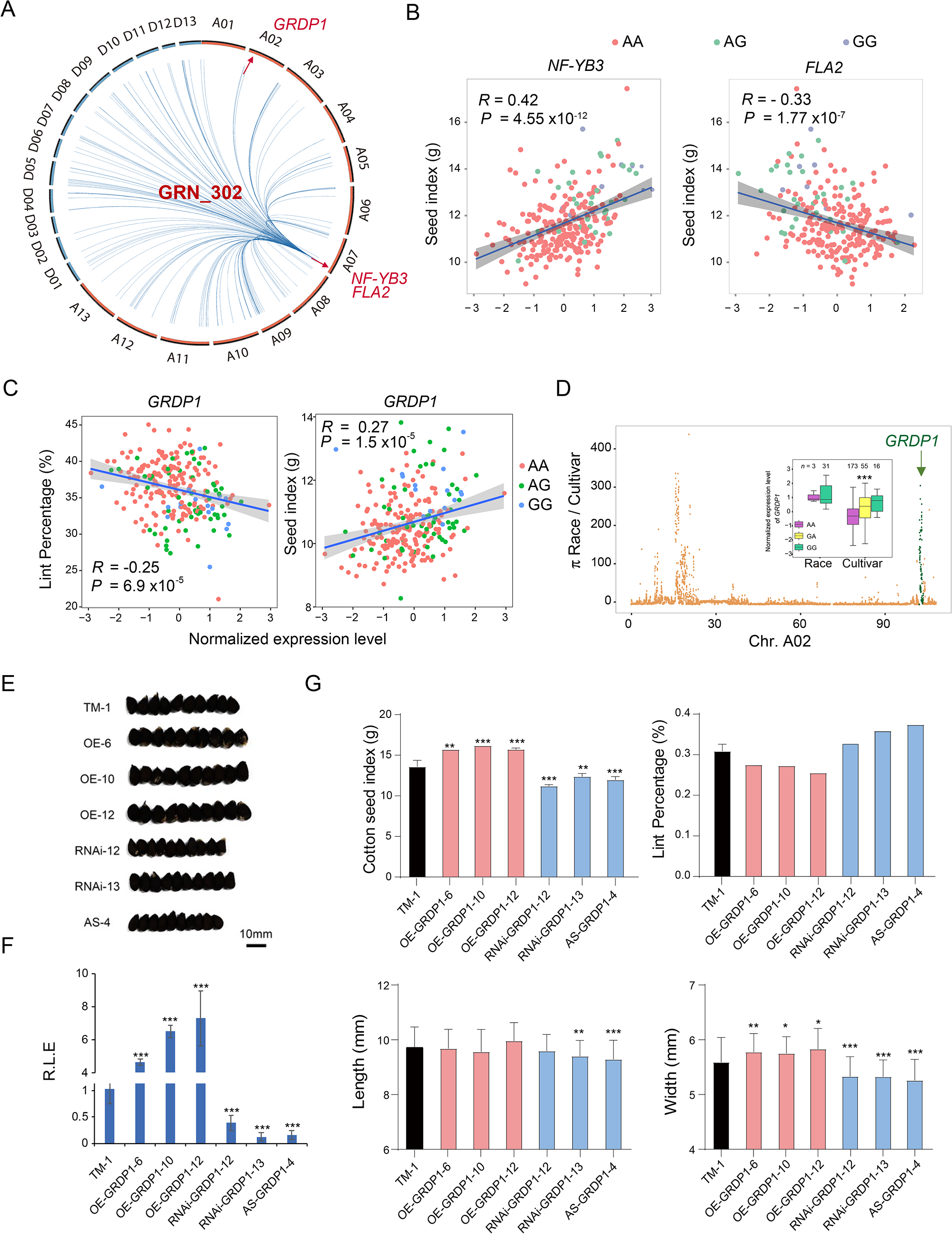
Validation of *GRDP1* as a candidate gene of seed size regulation. (A) Circos plot of autosomes indicating the association of eGenes with the locus underlying GRN_302 on Chromosome A07 (SNP A07:89225810). Lines with arrows indicate regulation of the expression of downstream genes. (B) Linear regression analysis of *FLA2* and *NF-YB3* expression and seed index phenotype; the scatter plot illustrates the high inter-tissue correlation. Each point represents one accession. Gene expression values were normalized by normal quantile transform. Points are colored based on SNP (A07:89225810) genotype. (C) Linear regression analysis of *GRDP1* expression and seed index. The scatter plot illustrates the high inter-tissue correlation. Each point represents one accession. Gene expression values were normalized by normal quantile transform. (D) Domestication sweeps in wild/landrace and cultivar populations. The value of π (wild) /π (cultivar) is plotted against position on chromosome A02. SNPs close to *GRDP1* are colored green. Box plot presenting differences of expression in each accession according to index SNP genotypes in the wild/landrace and cultivar populations. Boxes show the medians and interquartile ranges (IQR). The end of the top line is the maximum or the third quartile (Q) + 1.5× IQR. The end of the bottom line denotes either the minimum or the first Q − 1.5× IQR. Dots are either more than the third Q + 1.5× IQR or less than the first Q − 1.5× IQR. (*** *P* < 0.001, two-tailed Mann-Whitney test). (E) Photo image shows the seed size difference of wild type, *GRDP1-OE*, *GRDP1-RNAi* and *GRDP1-AS* lines. Bar, 10 mm. (F) The barplot shows the relative expression levels of wild type, *GRDP1-OE*, *GRDP1-RNAi* and *GRDP1-AS* lines (*** *P* < 0.001, two-tailed t test). (G) The histogram shows the weight of seed index, lint percentage, seed length and width from wild type, *GRDP1-OE*, *GRDP1-RNAi* and *GRDP1-AS lines*. (*** *P* < 0.001, two-tailed t test).

*GRDP1* is a *trans*-eGene in GRN_302, but a *cis*-eQTL in GRN_96(**Figures 4C and 7A**). The *GRDP1* homolog in *Arabidopsis, AtGRDP1* (*AT2G22660*), is predominantly expressed in the embryo during seed development and is involved with seed germination and ABA response ^58, 59^. Cultivars of the AA haploid type, which predominated in CUCP1, exhibited relatively low *GRDP1* expression (*P* < 10^-16^, Mann-Whitney test) (**Figure 7C**), and variation in *GRDP1* expression was found to be positively correlated with LP and negatively with SI (**Figure 7C**). In addition, *GRDP1* in GRN_96 is located in a selective sweep region associated with domestication (**Figure 7D**) and exhibits fixed haplotypes in the cultivated population within CUCP1 (**Figure 7D**). The inclusion of *GRDP1* in a selective sweep region was also reported in a latest study using an Upland cotton population with a large sample size ^60^.

To confirm the function of *GRDP1*, we constructed transgenic cotton to alter the expression of the endogenous *GRDP1*. The T_2_ generation of transgenic seeds from multiple independent lines of overexpressing *GRDP1 (OE-GRDP1)*, RNAi *(RNAi-GRDP1)* and antisense of *GRDP1* (*AS-GRDP1)* were obtained (**Figure 7E**). Real-time PCR examination confirmed that *GRDP1* expression were significantly higher in the *OE-GRDP1* lines than in the wild-type (WT), and significantly lower in the RNAi lines and antisense lines (**Figure 7F**). Furthermore, the seed index was significantly larger in the *OE-GRDP1* lines than in the WT lines, while it was significantly lower in *AS-GRDP1* and *RNAi-GRDP1* lines (**Figure 7G**). The seed width and length showed the a similar trend. On the contrary, LP was higher in *AS-GRDP1* and *RNAi-GRDP1* lines while it was significantly higher in *OE-GRDP1* lines compared to WT (**Figure 7G**).

The above data confirmed that *GRDP1* has a direct effect on cotton seed size, and the natural variations in *GRDP1* structure and expression are of great potential in improving seed cotton yield in cultivated cotton populations. Moreover, the top-ranked eGenes in the integrative study are demonstrated to be the causal genes in the seed size associated GWAS loci.

## Discussion

### Mining potentially functional genes using an eQTL map

In this study, the eQTL map of 1-DPA ovule was constructed, comprising 12,207 eQTLs. This map represents a bridge between phenotype-associated variation and gene expression. In total, 66 out of 187 reported phenotypic GWAS loci were colocalized with *cis-* and *trans-*eQTLs.

Most studies prioritizing genes for complex traits have considered only *cis*-eQTL effects, despite the fact that *trans*-eQTLs account for a substantial portion of the eQTL regulation network. *Trans*-eQTLs are routinely ignored because their effects are in general as weak as each individual variation ^38^ and are considered to be more tissue-specific than *cis*-eQTLs ^61^. In this study, we estimated heritability based on the variants of both *cis*- and *trans-*eGenes and successfully captured the missing heritability of seed size related traits (**Figure 6**), which emphasizes the importance of *trans-*eGenes.

Crop domestication is often associated with genomic sweeps, which rid cultivated populations of rare variants ^11^. Thus, rare variants associated with phenotypes are difficult to detect by GWAS in a cultivated population ^62^. However, if any critical genetic variants are present in an upstream regulatory module, the associated variations in important functional genes should be noticeable accordingly. The effects of these genetic variants on downstream gene expression in a GRN can also be detected as *trans*-eQTLs. The presented results demonstrate that integrative analysis using eQTL and GWAS to construct a GRN can retrieve part of the “lost heritability” by mining a comprehensive resource that points to impactful genes.

### Crosstalk over GRNs reveals the pleiotropy of GWAS loci

Most agronomic traits are controlled by multiple quantitative loci, and many phenotypes tend to be integrated or controlled by pleiotropic genes ^63^. Interaction between loci, i.e. epistasis, also increases the complexity of the genetic basis of a phenotype. Thus, the dissection of a single gene or a gene related to one specific trait is insufficient for molecular breeding.

In the present study, we found that different genetic variants associated with related but different phenotypes can influence the expression of the same genes via *cis*-or *trans*-regulatory mechanisms. To identify the eGenes that are coordinately affected by different loci, independent functional GRNs were compared in a pairwise manner, which revealed many *cis-*eGenes to be shared across GRNs as *trans-*eGenes. This crosstalk between GRNs via *trans*-regulation by eGenes provides insight into the pleiotropy of GWAS loci. Ultiizing these pleiotropic genes from GWAS loci can aid in in engineering multiple desirable phenotypes through design.

### Prioritizing Impactful genes using machine learning

Machine learning methodologies are promising for the analysis of complex biological data. When implementing such methodologies, having a large p and small n poses a major challenge ^64^. In particular, the large number of genomic variations and genes with expression data that need to be examined can dramatically increase the computational cost, especially since they are usually much more numerous than the samples ^65, 66^. This study demonstrates that using genes colocalized in GWAS loci and eQTL GRNs can effectively reduce the dimensionality of biological data for machine learning. Specifically, it narrows the focus from 49,637 expressed genes down to 701 trait-associated genes. By employing XGBoost training, the impactful ranking of the 661 genes in cotton yield GRNs was obtained. This strategy of GRN navigation successfully identified the top-ranking genes in seed size regulation.

In addition to eQTLs, a wide spectrum of advanced molecular biotechnologies can be used to construct GRNs and reduce the dimensionality of biometrics for machine learning. Typically, GRNs can be characterized based on physical interactions and upstream and downstream regulatory relationships ^67^. Among the state-of-the-art methods for GRN construction, one option is the three-dimensional genome (3D genome), which includes extensive DNA-DNA and DNA-RNA interactions ^68^. Single-cell multi-omics can also be applied to uncover GRNs ^69^. In future study, GRNs identified through different platforms and technologies can be further utilized for machine learning to prioritize impactful genes.

## Materials and methods

### Plant material and growth conditions

A total of 279 accessions were collected from the Institute of Cotton Research at CAAS, including 34 wild/landrace *Gossypium hirsutum* (*Gh*) accessions, such as *G. palmeri*, *G. punctatum*, *G. morrilli*, *G. yucatanense*, *G. richmondi*, *G. marie-galante*, and *G. latifolium*, as well as 245 core germplasm samples (**Table S1**). The core germplasm accessions were previously genotyped by our laboratory ^12^, while the whole-genome sequencing of 34 wild accessions was newly conducted in this study (**Table S2**). Plants of the 279 accessions were grown in a farm environment during the summer of 2018 in Dangtu, Anhui, China. Two independent biological samples were taken from each accession and grown in different experimental fields. For ovule collection, 16-18 plants were grown for each accession; the 1-DPA ovules collected were then bulked for total RNA extraction and sequencing. Leaves from the 34 wild/landrace *Gh* accessions were collected for DNA extraction, sequencing, and genotyping.

Phenotypic data for nine complex traits (seed index [SI], boll weight [BW], boll number [BN], lint percentage [LP], fiber elongation [FE], fiber micronaire, fiber length [FL], and fiber strength [FS]) were collected over three years (2007, 2008, and 2009) from nine environments: three farms each in Anyang (AY) in the Yellow River cotton-growing area, Nanjing (NJ) in the Yangtze River cotton-growing area, and Korla in Xinjiang (XJ), the northwestern cotton-growing area ^12^. The best linear unbiased prediction (BLUP) values (Bates et al., 2015) were estimated for different phenotypes using the R package lme4. These values segregated the genetic and environmental effects that influence the phenotypes ^70^.

### Sample preparation

Genomic DNA of 34 wild/landrace *Gh* accessions was extracted from young leaves using the CTAB method. For RNA profiling, 1-DPA ovules were harvested from 12:00 to 1:00 pm. The aim was to collect samples in the shortest amount of time possible so as to minimize the effects of physiological changes. Harvested ovules were frozen with liquid nitrogen for RNA extraction. Total RNA was extracted using the Trizol method, following the the manufacturer’s instructions, and RNA quality was verified with an Agilent 2100 Bioanalyzer. Transcriptome libraries were constructed using the standard Illumina RNA-seq protocol (Illumina, Inc., San Diego, CA, catalog no. RS-100-0801). RNA and DNA sequences were generated as 150 bp paired-end reads from libraries with inserts of 350 bp.

### SNP identification and annotation

WGS data were quality controlled using fastp (V 0.12.2) with default parameters. ^71^. Genome and annotation files of TM-1 v2.1 ^36^ were indexed using BWA index with the flag (-a bwtsw) ^72^. Reads were mapped to the reference genome using the BWA. SAM files were sorted, indexed, and converted to BAM files using SAMtools (V 1.16) . ^73^. Only uniquely mapped non-duplicated reads were used for SNP calling according to the best practices pipeline of GATK (v3.7)^74^. Duplicated reads in the resulting alignment BAM files were marked using Picard Tools (http://picard.sourceforge.net). SNPs were called based on a minimum phred-scaled confidence threshold of 20 (-stand_call_conf >20) using the GATK tool HaplotypeCaller and then filtered using the GATK tool VariantFiltration with the following requirements: Fisher strand value (FS) < 30.0 and quality by depth value (QD) > 2.0. For GWAS and eQTL analysis, SNPs having a high missingness rate (> 20%) or low minor allele frequency (MAF < 0.05) were removed using VCFtools (V 0.1.13) with the parameters (--remove-indels, --maf 0.05, --max-maf 0.95, --max-missing 0.8) ^75^. Missing genotypes were imputed using Beagle with the parameters (window = 50000, overlap = 5000, ibd = True)^76^. This process identified ∼1.19 million autosomal SNPs, output in a variant call format (VCF) file.

### LncRNA annotation

To examine the expression of non-coding sequences, we performed population-level transcript assembly of long non-coding RNAs. RNA-seq data were quality controlled using fastp (V 0.12.2) with default parameters (Chen et al., 2018). An average of 24.34 million reads was obtained for each library. Clean RNA-seq reads (150 bp paired-end) were aligned to the *Gh* TM-1 v2.1 reference genome using Hisat2 (V 2.1.0) with parameter (--dta) ^77^. Mapped reads in each library were subsequently passed to StringTie (V 2.0) for transcript assembly ^77^ using annotated TM-1 transcripts ^36^ as a reference transcriptome; the transcripts so assembled were combined into a unified set using cuffmerge with parameter (-c 3) ^78^. Transcripts of less than 200 nt were discarded. Using Cuffcompare (V 2.2.1), transcripts were given a class code of “u,” “x,” or “i,” respectively representing intergenic sequences, antisense sequences of known genes, and intronic sequences. The Coding Potential Calculator2 (CPC2) (V 0.1) was used to calculate the coding potential of transcripts of each given class (“u,” “x,” or “i”) with default parameters. All transcripts with CPC scores > 0 were discarded. The remaining transcripts were subjected to pfam_scan in order to exclude those containing known protein domains (cutoff < 0.001) ^79^. The transcripts left after that step were considered candidate lncRNAs. To reduce isoform complexity, only the longest transcript of each locus was used for further analysis.

### Expression profiling

Gene expression of the newly annotated transcripts, including lncRNAs, was quantified using StringTie (V 2.0) ^77^. Pearson’s correlation coefficient was calculated for replicates using the cor () function in R. For comparison of transcriptomes across different tissues, raw RNA-seq were analyzed through our bioinformatics pipeline as described above ^44^. Heatmaps of the expression of eGenes belonging to GRN_302 and GRN_808 were generated using the *pheatmap* package (https://cran.r-project.org/web/packages/pheatmap).

### Genome-wide association analysis of eQTLs

The analysis included 279 individuals for whom genotype and gene expression data were available. GWAS was performed for those accessions with a total of 1.19 million SNPs (MAF > 5% and missing rate < 20%). Population structure was calculated using GCTA (V 1.92.1) with the parameters (--make-grm --pca) ^48^. Only genes having FPKM > 1 in more than 5% of accessions were defined as expressed for the purpose of eQTL mapping. The expression of each gene was normalized using QQ-normal in R as is commonly done in QTL studies ^80^. Ultimately, a dataset comprising 42,858 PCGs and 6,779 lncRNAs was obtained and used to conduct downstream analyses. The first three genotyping principal components (PCs) and kinship matrix were employed as covariates to control false-positive associations. Genotype files were transposed using plink (V 1.9) with the parameters (--bfile –recode12 –output-missing-genotype0 – transpose --out) ^81^. Kinship matrices were obtained using the emmax-kin function of EMMAX with parameters (-v -d 10) ^37^. eQTL mapping was carried out using EMMAX with a mixed linear model and parameters (-v -d 10 -t -o -k -c) ^37^. The effective number of independent SNPs was calculated using the Genetic type 1 Error Calculator (GEC), and significant SNPs were identified using the threshold of *P* < 2.18 × 10^-6^ suggested by GEC ^82^.

To reduce eQTL redundancy, we conducted linkage disequilibrium (LD) analysis for the associated SNPs. Lead SNPs within the LD block (*R*^2^ > 0.1) for each trait were merged into one eQTL using plink (V 1.90) with parameters (-r2 -l -window 99999) ^81^. The eQTLs were then further classified as *cis* or *trans* based on the distance between the marker SNP and the transcription start sites or transcription end sites of associated genes (threshold: 1 Mb)^38^. Hotspots were identified using hot_scan with parameters (-m 5000, -s 0,05) ^83^. *Cis-* and *trans-*eGenes in GRN_302 were visualized using Cytoscape (version 3.4.0; www.cytoscape.org) ^84^.

### Construction of GRNs

Linkage disequilibrium (LD) pruning was performed to provide a list of independent GWAS variants for downstream analyses. Pruning was carried out according to three linkage disequilibrium thresholds (*R*^2^ > 0.1) using an in-house Perl script. To test whether the eGenes within a GRN have more connections and interactions among themselves, we downloaded PPI pairs from the STRING database (https://stringdb-static.org/download/protein.links.v11.5/3702.protein.links.v11.5.txt.gz) ^46, 85^, which consisted of 16,029,730 PPI pairs.

### Gene function enrichment analysis

To determine whether genes within a GRN share common functional features, we performed GO term enrichment analyses using a hypergeometric test in GOstats (V 2.50.0) ^86^. GO terms were retrieved from the annotation files of TM-1 ^36^, and categories that contained at least five genes were considered significantly enriched if having a false discovery rate-corrected *P* < 0.05.

### GRN effect on heritability

Two GWAS catalogs previously published were employed to assess the impactsof SNPs in GRN contribute to phenotypic variability ^12, 87^. The analysis considered six complex agronomic traits: seed index (SI), boll weight (BW), boll number (BN), lint percentage (LP), fiber strength (FS), and fiber length (FL). The association of phenotypic variation with the GRN was evaluated using Genomic-Relationship-matrix Restricted Maximum Likelihood (GREML), performed in GCTA ^48^. Three datasets were produced: (1) SNPs from *cis*/*trans* eGenes (n = 216) within GRN, (2) eSNPs in GRN_302 and randomly selected eGenes not in GRN, and (3) SNPs from randomly selected genes. The test and control groups in all three datasets used the same number of SNPs. The genetic relationship matrices for those datasets were built using GCTA (v 1.92.1) with the parameter (make-grm), then estimated the amount of phenotypic variation in FL, FS, FU, and LP that could be explained by each SNP set using GCTA with the parameter (mgrm) ^48^. We repeated this process 100 times, each time randomly sampling the set of SNPs.

### Machine learning models for trait prediction

The predictive model for phenotype based on gene expression was constructed using an ensemble of gradient boosted trees (XGBoost) ^88^. The eGenes belonging to pSNP-eGene pairs were considered to be informative genes. For the SI trait, the 246 available individuals were initially partitioned into training and testing datasets consisting of 90% and 10% of the data, respectively. The testing samples were never used in training.

For prediction, we applied the XGBoost ^88^ module of python. The XGBoost classifier is a gradient boosting method. The goal function of the XGBoost algorithm model is *obj* (*θ*) = *L*(*θ*) + Ω (*θ*), where *L*(*θ*) is the training loss function and Ω(*θ*) is the complexity function of the tree. 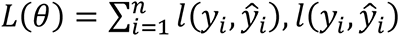 corresponds to the training loss function for each sample, where *y*_*i*_ represents the true value of the *i*th sample and *ŷ_i_* represents the estimated value of the *i*th sample. Then, 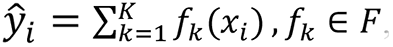, where *K* represents the number of trees, *F* represents all possible *DT*, and *f* denotes a specific CART tree. 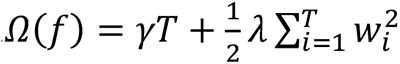, in which *w_i_* score on the *i*th leaf node and *T* is the number of leaf nodes in the tree. By adjusting the parameters, the objective function was continuously optimized, and optimal results were ultimately obtained ^24^. The grid search algorithm was used to optimize hyper-parameters in each iteration, which included max_depth, min_child_weight, gamma, subsample, col-sample_bytree, and learning_rate.

This process was repeated 100 times using different seeds to take into account the variation in the hyperparameter optimization procession. As a description of stability of the individual phenotype predictions, we computed the mean square error (MSE) and *R*^2^ of the predictions in the test set. Finally, the importance of each gene was calculated.

### Transcriptome-wide association (TWAS)

The TWAS was carried out using the functional summary-based imputation (FUSION) approach (http://gusevlab.org/projects/fusion/) ^89^. This method precomputes the functional weights of gene expression, and then integrated them with summary-level GWAS results to impute the association statistics between gene expression and phenotype. The FUSION approach only considers *cis*-eGenes, typically within 500 kb or 1 Mb; in this work, 1,085 cis-eGenes were included in the analysis.

### Transgenic cotton and *Arabidopsis*

The transgenic cotton was transformed by WIMI Biotechnology Co., Ltd. using a shoot apical meristem (SAM) cells-mediated transformation system (SAMT) ^90^. To construct the over-expression and suppression vectors of *GRDP1*, total RNA was isolated from the cotton TM-1 using Trizol reagent (Invitrogen) according to the manufacturer’s instructions. And was then treated with DNase I (Promega). First-strand cDNA was then synthesized using M-MLV reverse transcriptase (Promega). The Open read frames (ORFs) of *GRDP1* were amplified by regular PCR with added *Xba*I and *Bam*HI, and then inserted into the basic vector pWMV062-AADA controlled by the constitutive Cauliflower mosaicvirus (CaMV) 35S promoter. *OE-GRDP1* and *AS-GRDP1* constructs were introduced into *G. hirsutum* accession TM-1 via *Agrobacterium tumefaciens* (strain LBA4404) using SAMT ^90^. The T_2_ homozygous transgenic lines (confirmed by target gene PCR, target protein detection, and target gene real-time qPCR) were used for further analysis. The primers used for vector construction and PCR-based screening are provided in **Table S11.**

To generate transgenic over-expression lines of *Arabidopsis* plants, the coding region of *NF-YB3* were cloned into *Xba* I and *Bam*H I restriction sites of *pBI121* binary vectors, under the control of the CaMV 35S. The *pBI121*_*NF-YB3* plasmids were transformed into *Arabidopsis thaliana* Col-0 by *A. tumefaciens* (GV3101) using the floral dip method. Primers are listed in **Table S11**. The integration of the transgene into different transgenic lines was confirmed by PCR.

## Data availability

All RNA and DNA sequencing reads have been deposited in the NCBI Short Read Archive (https://www.ncbi.nlm.nih.gov/sra) under Bioproject PRJNA730082. Sample IDs and metadata can be found in Supplementary Data 1.

## Acknowledgements

This work was financially supported in part by grants from the National Key Research and Development Program (2022YFF1001400), National Natural Science Foundation of China (NSFC 3200379) and Fundamental Research Funds for the Central Universities and JCIC-MCP. We thank the Chinese national medium-term cotton gene bank at the Institute of Cotton Research (ICR) of the Chinese Academy of Agricultural Sciences (CAAS) and National Wild Cotton Nursery, Sanya, China for kindly sharing the cotton accessions.

## Author contributions

X.G. conceptualized the project. L.W., M.H., J.H., S.W., K.N., M.L., M.G., Z.C., H.Z., K.W., and J.L. conducted the experiments. T.Z. performed the bioinformatics analysis. T.Z., Z.T., and X.G. prepared the manuscript. All authors read and approved the final manuscript.

