## Supplementary material for "Integration of eQTL and Machine Learning Methods to Dissect Causal Genes with Pleiotropic effects in Genetic Regulation Networks of Seed Cotton Yield": Table S1-Table S14

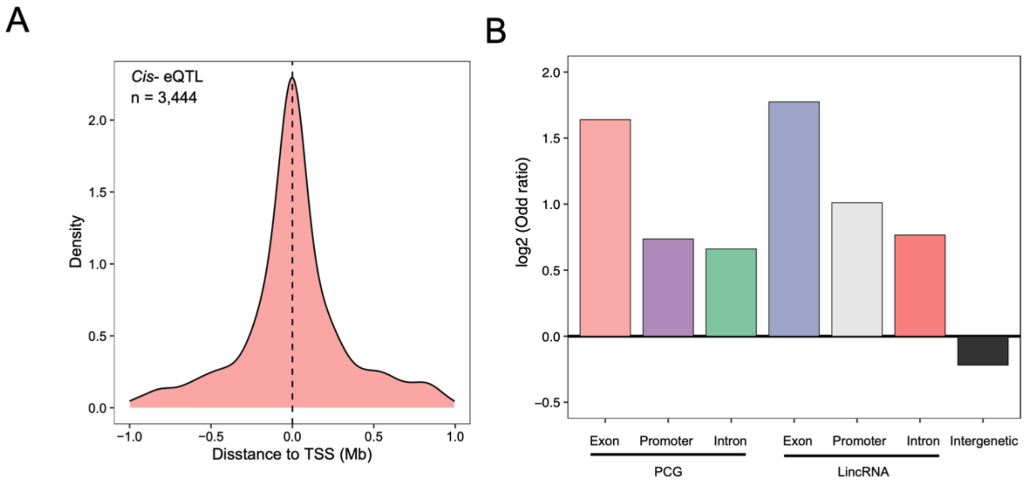


**Figure S1: Genomic feature of eQTL**

(A) Distribution of distance of eQTL to transcription start or termination sites (TSS, TTS) of their associated eGene in a ± 1000 kb window, shown for eQTLs in linkage disequilibrium. (B) Enrichment of eSNP overlapping genomic features. The proximal promotor is defined as 2000 bp upstream of the transcription site.

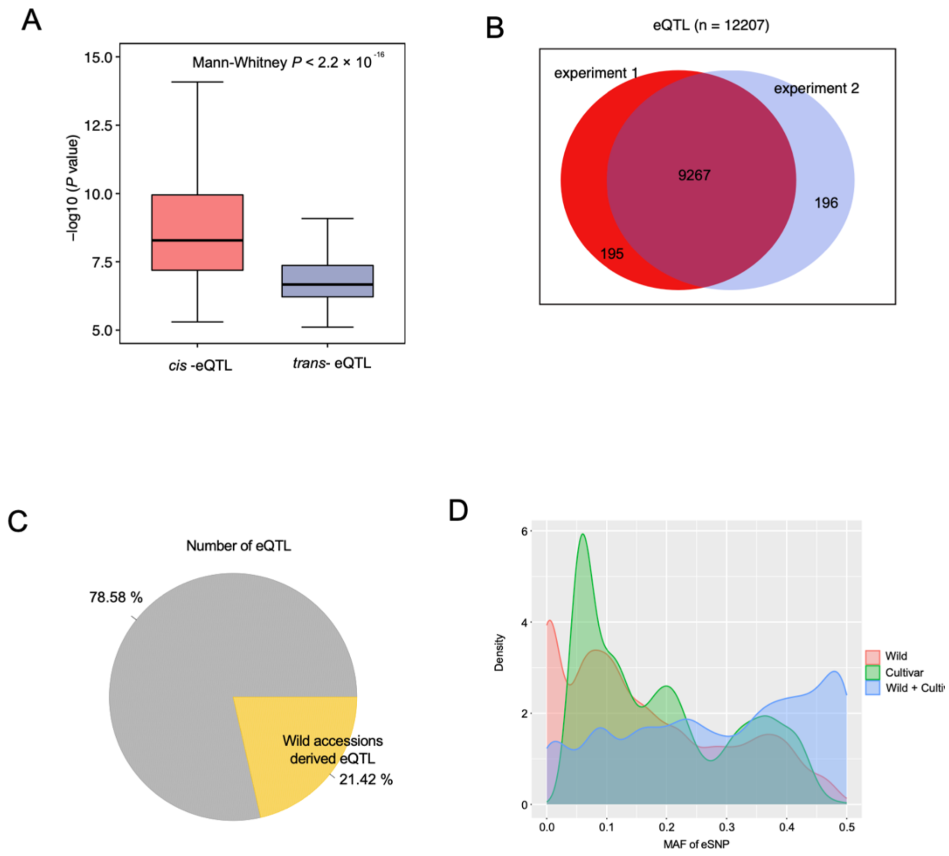


**Figure S2: Reliability assessments of eQTLs.** (A) Box plot showing the distribution of -log10 (*P value*) of *cis*- and *trans*- eQTL. Boxes show the medians and IQRs. The end of the top line is the maximum or the third quartile (Q) + 1.5× IQR. The end of the bottom line denotes either the minimum or the first Q − 1.5× IQR. The dots are either more than the third Q + 1.5× IQR or less than the first Q − 1.5× IQR. The *P value* between *cis*-eQTL and *trans*-eQTL were calculated based Mann-Whitney test. (B) Veen Plot showed the eQTL supported by experiment 1 and experiment 2. (C) Pie plot showed the number of eQTL showed the difference between wild and cultivar accessions. (D) MAF of the eSNP in wild and cultivar populations.

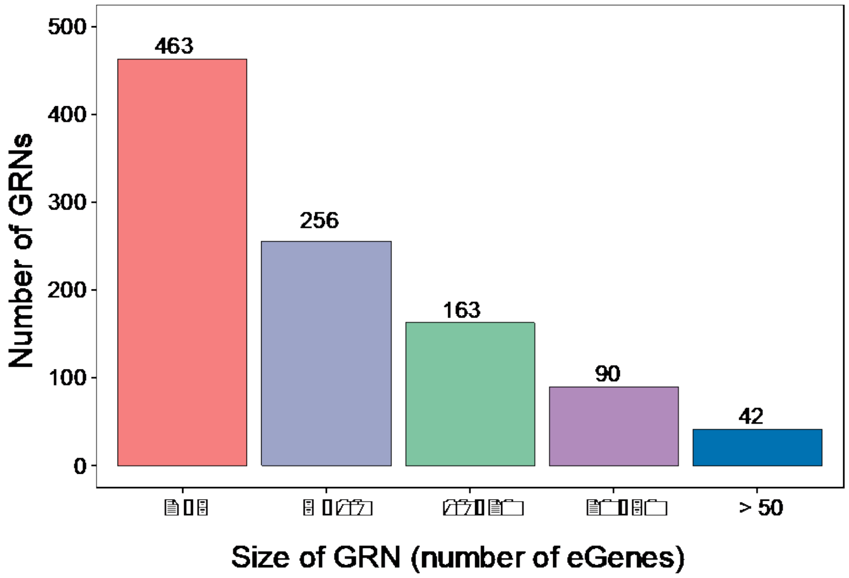


**Figure S3: Bar plot showing the numbers of GRNS with different size ranges.**
X axis represents the size ranges (number of genes in each GRN). Y axis represents the number of GRN.

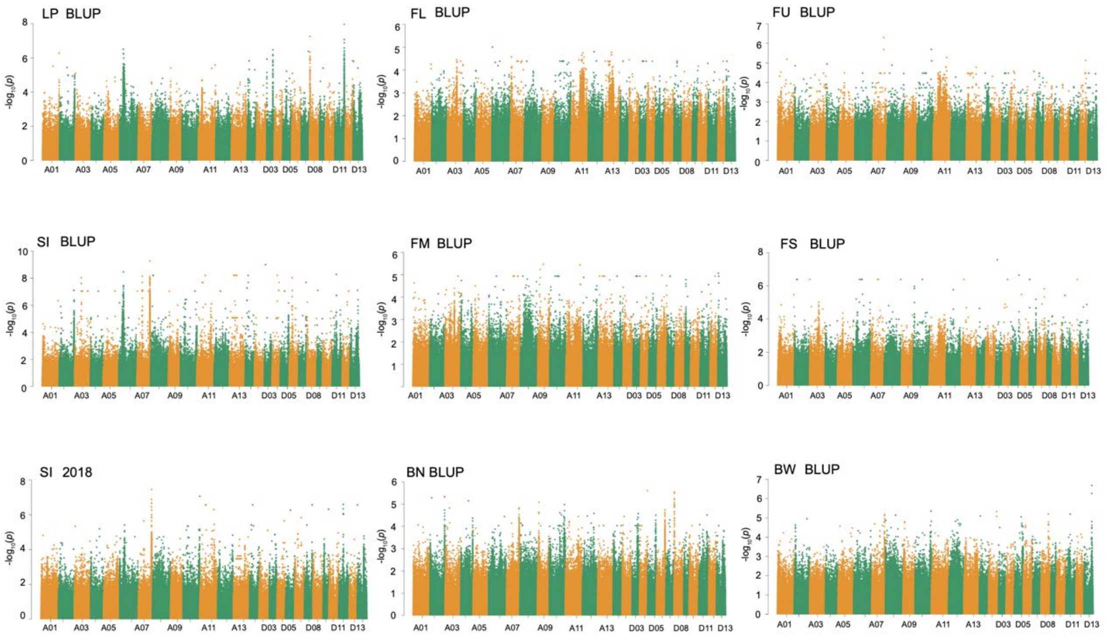


**Figure S4: Manhattan plots for different traits.** The traits were calculated based on BLUP values.

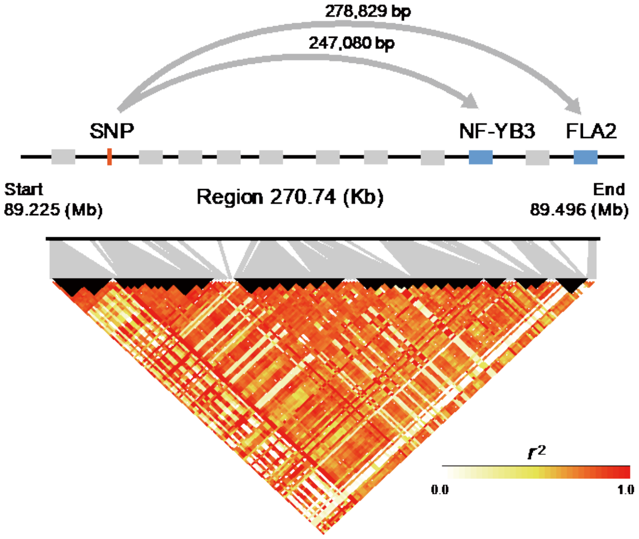


**Figure S5: Locus Zoom plot (Chromosome A07: from 89.23 to 89.50 Mb) showing that the lead SNP locus for SI and BW is in linkage disequilibrium and colocalizes with an eQTL associated with *NF-YB3* and *FLA2*.** No other gene in the locus was implicated based on linkage disequilibrium and eQTL co-localization.

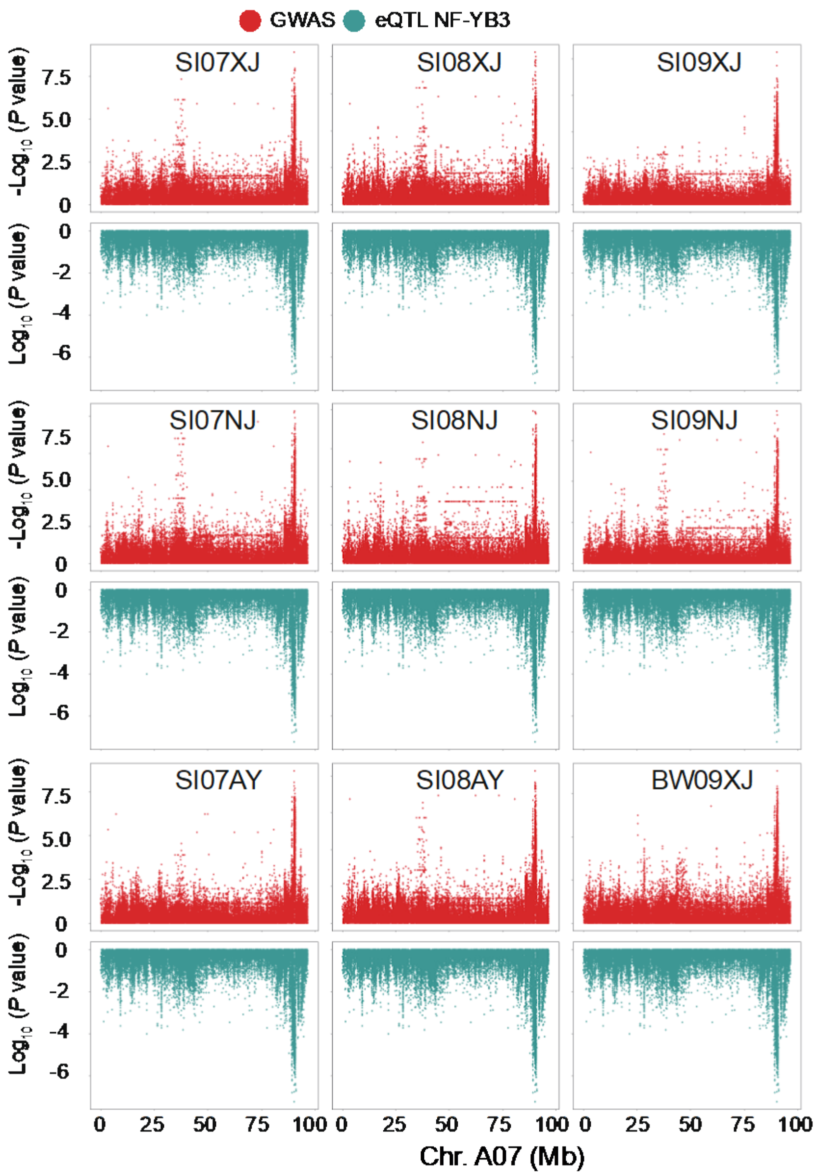


**Figure S6: Combined Manhattan plots for different traits and gene expression.** Manhattan plot for SI and BW traits (Top), expression of *NF-YB3* (bottom), respectively for three years and three locations. Genomic position is shown on the x-axis, and each SNP associated with the indicated trait on the y-axis. Points represent SNPs.

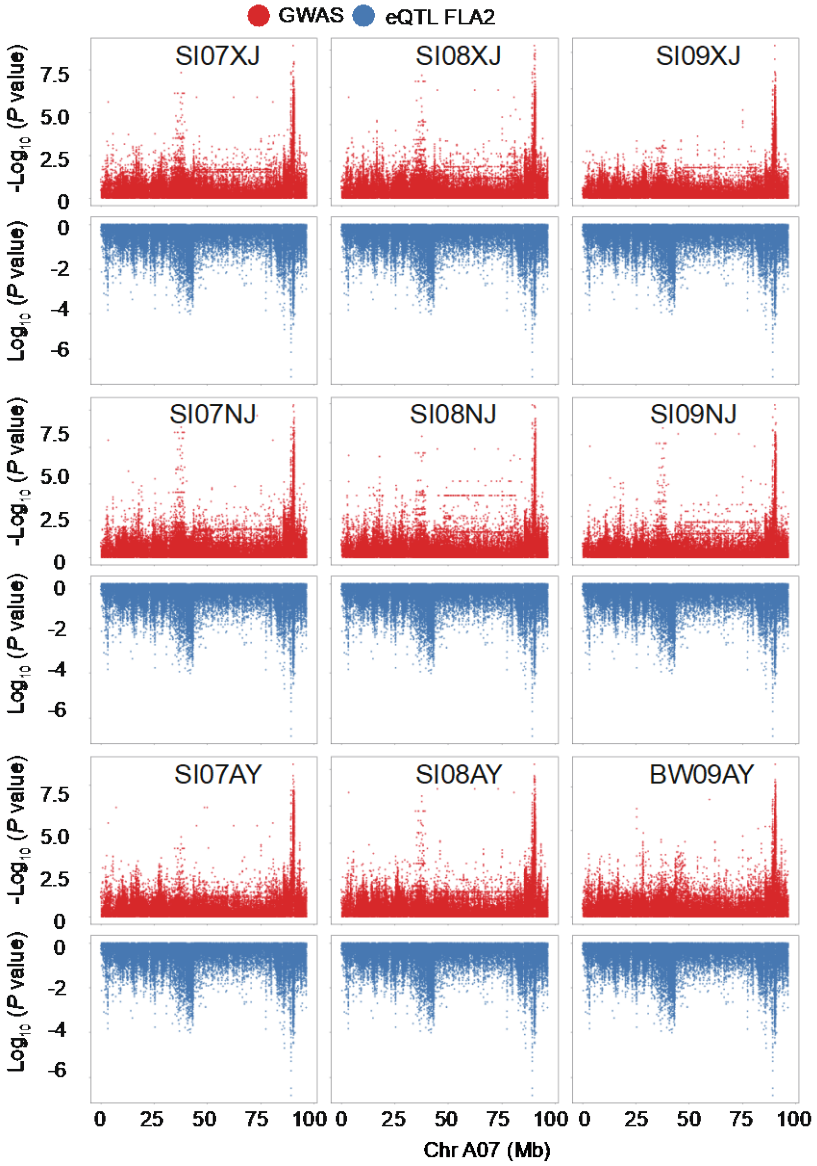


**Figure S7: Combined Manhattan plots for different traits.** Manhattan plot for SI, BW(Top), expression of *FLA2* for three years and three locations, respectively. Genomic position is shown on the x-axis, and each SNP associated with the indicated trait on the y-axis. Points represent SNPs.

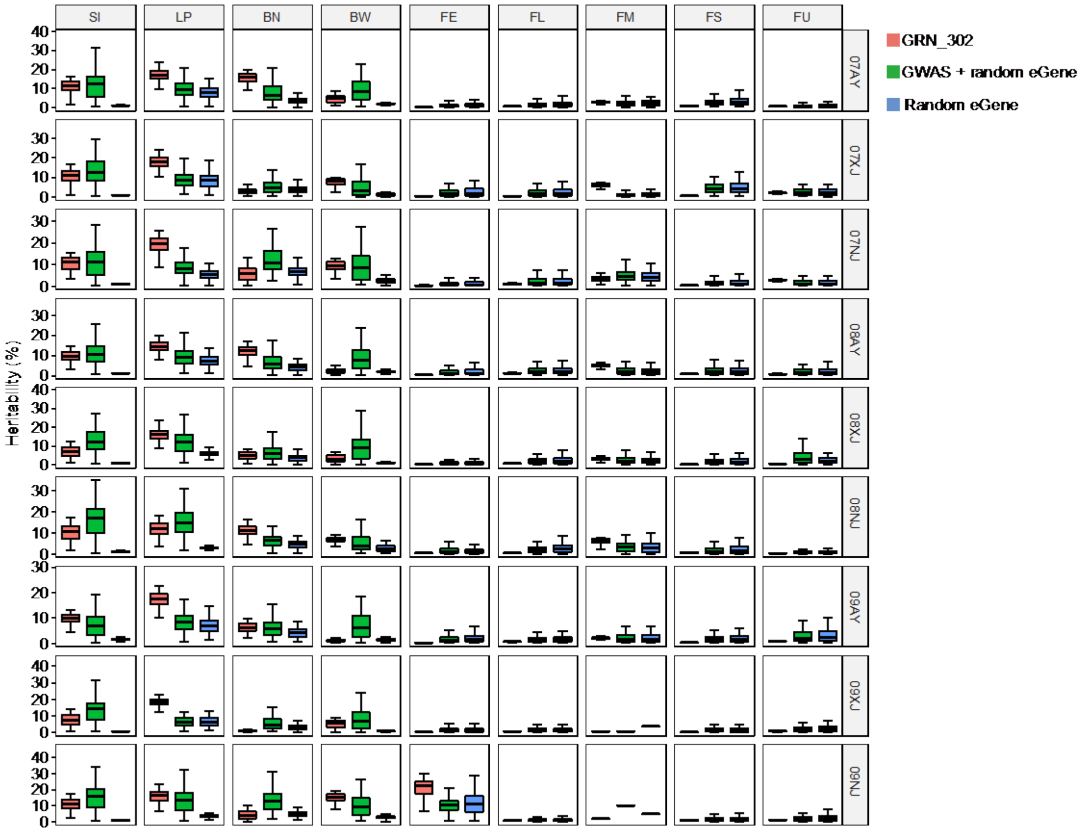


**Figure S8: GRNs_302 explained a larger fraction of heritability than GWAS loci.** Box plots show the estimated heritability ($h^2$) across different phenotypes explained by different sets of eSNPs, Boxes show the medians and IQRs. The end of the top line is the maximum or the third quartile (Q) + 1.5× IQR. The end of the bottom line denotes either the minimum or the first Q − 1.5× IQR. Dots indicate values outside those bounds.

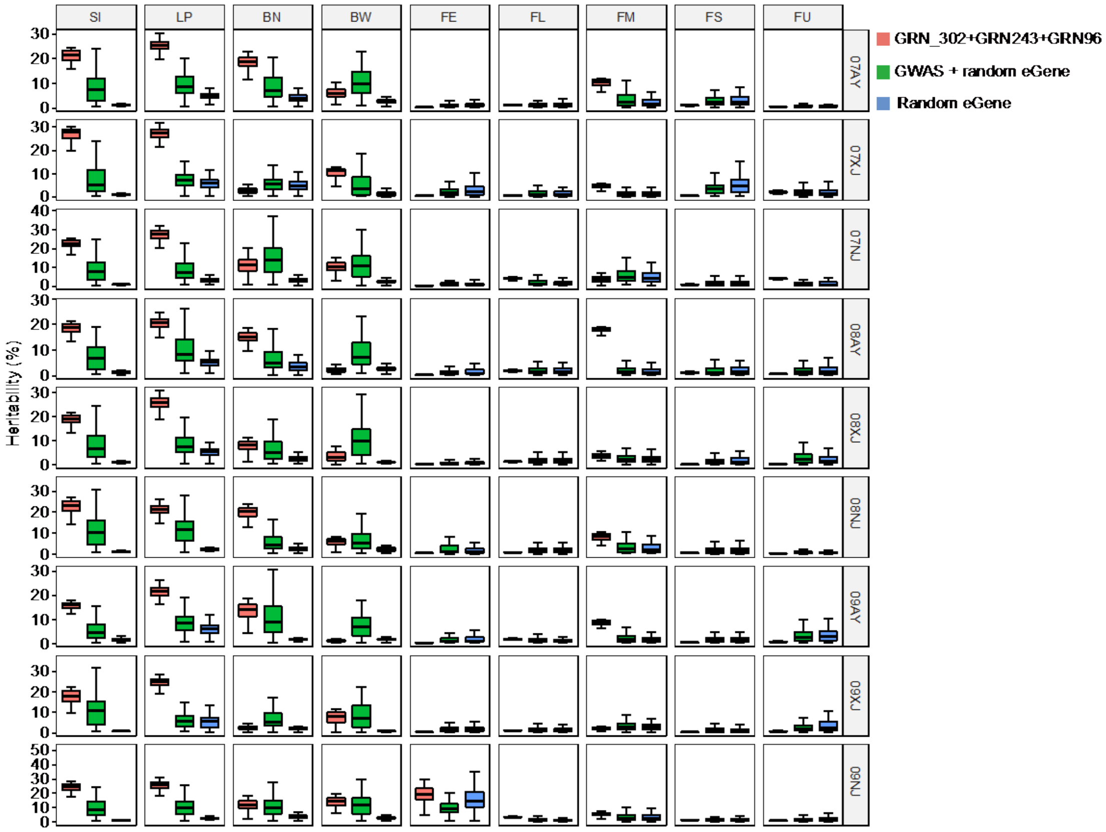


**Figure S9: GRNs_302+GRN_243+GRN_96 explained a larger fraction of heritability than GWAS loci.** Box plots show the estimated heritability ($h^2$) across different phenotypes explained by different sets of eSNPs, Boxes show the medians and IQRs. The end of the top line is the maximum or the third quartile (Q) + 1.5× IQR. The end of the bottom line denotes either the minimum or the first Q − 1.5× IQR. Dots indicate values outside those bounds.

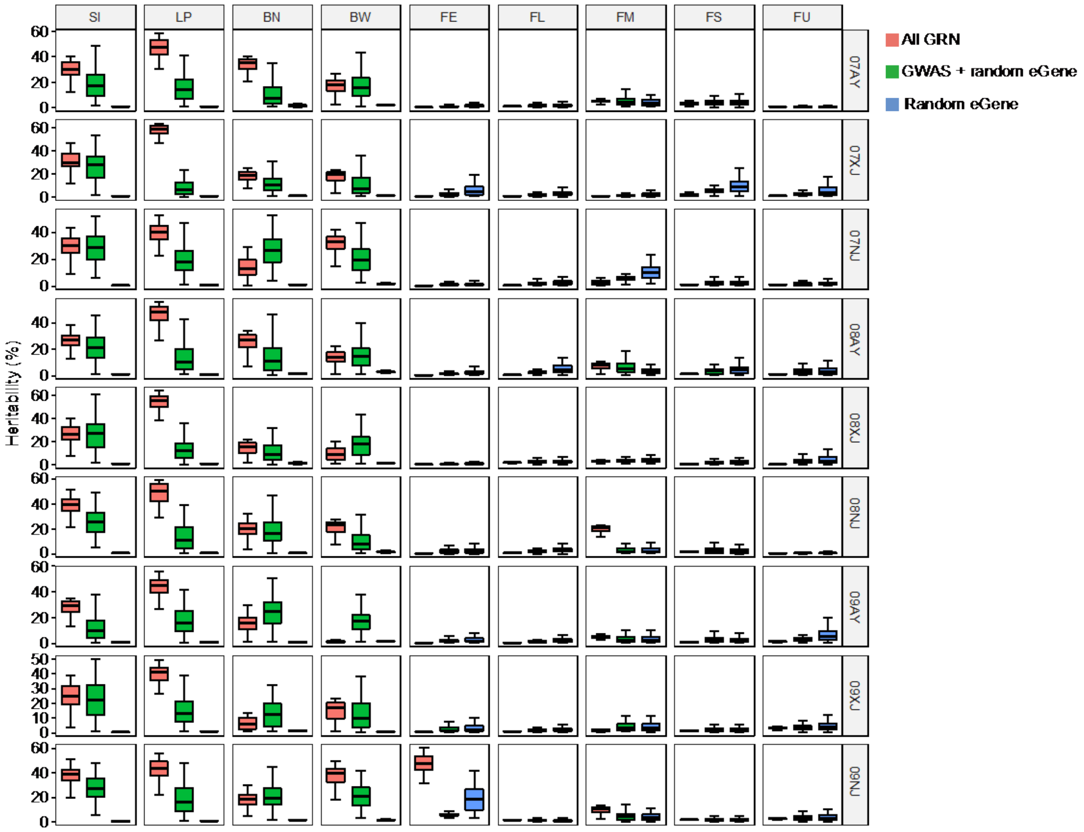


**Figure S10: GRNs explained a larger fraction of heritability than GWAS loci.** Box plots show the estimated heritability ($h^2$) across different phenotypes explained by different sets of eSNPs, Boxes show the medians and IQRs. The end of the top line is the maximum or the third quartile (Q) + 1.5× IQR. The end of the bottom line denotes either the minimum or the first Q − 1.5× IQR. Dots indicate values outside those bounds.

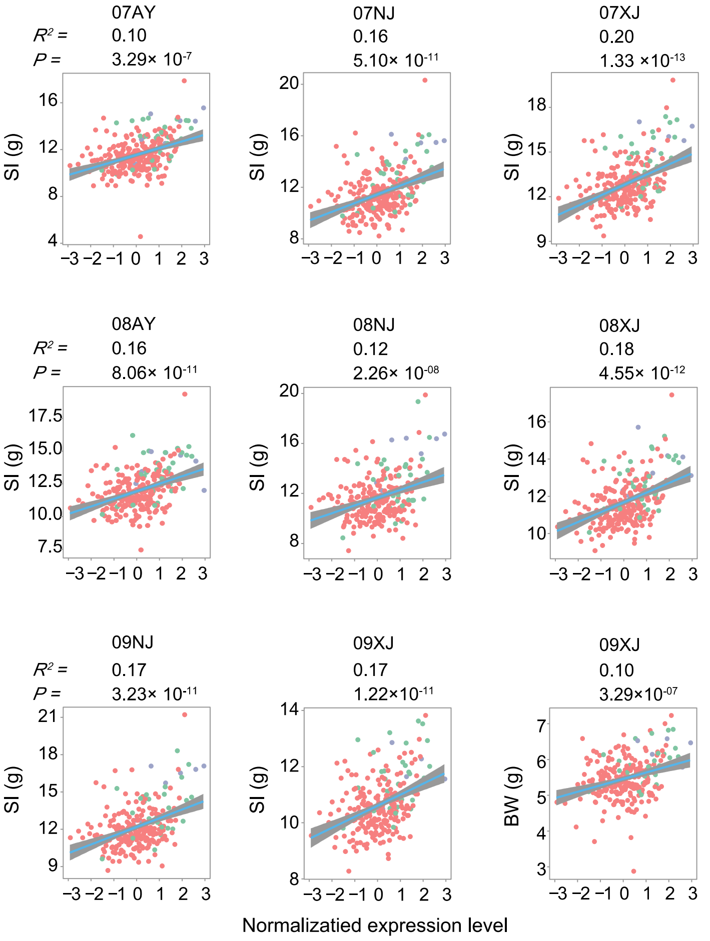


**Figure S11 Linear regression analysis show the correlation between *NF-YB3* expression and SI in the tested Upland cotton population.** The scatter plots illustrate the high inter-tissue correlation for three years and three locations. Each point represents one accession. Gene expression values were normalized by normal quantile transform.

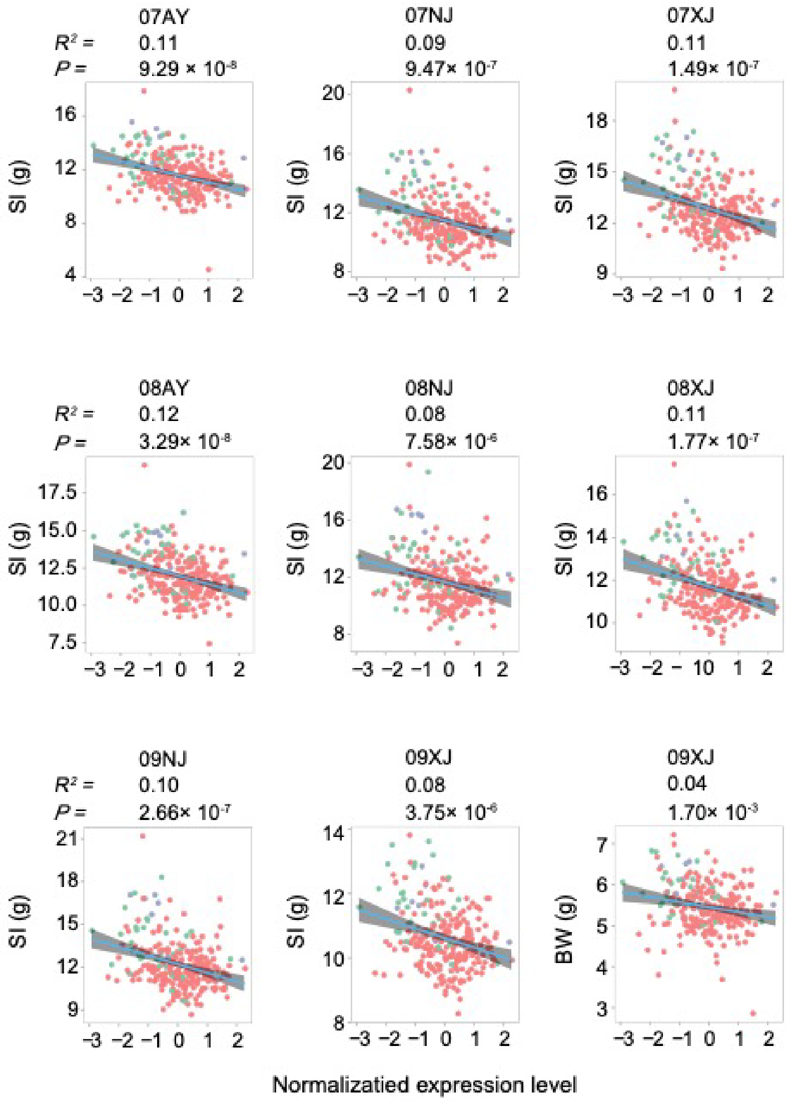


**Figure S12 Linear regression analysis show the correlation between *FLA2* expression and SI in the tested Upland cotton population.** The scatter plots illustrates the high inter-tissue correlation. Each point represents one accession for three years and three locations, respectively. Gene expression values were normalized by normal quantile transform.

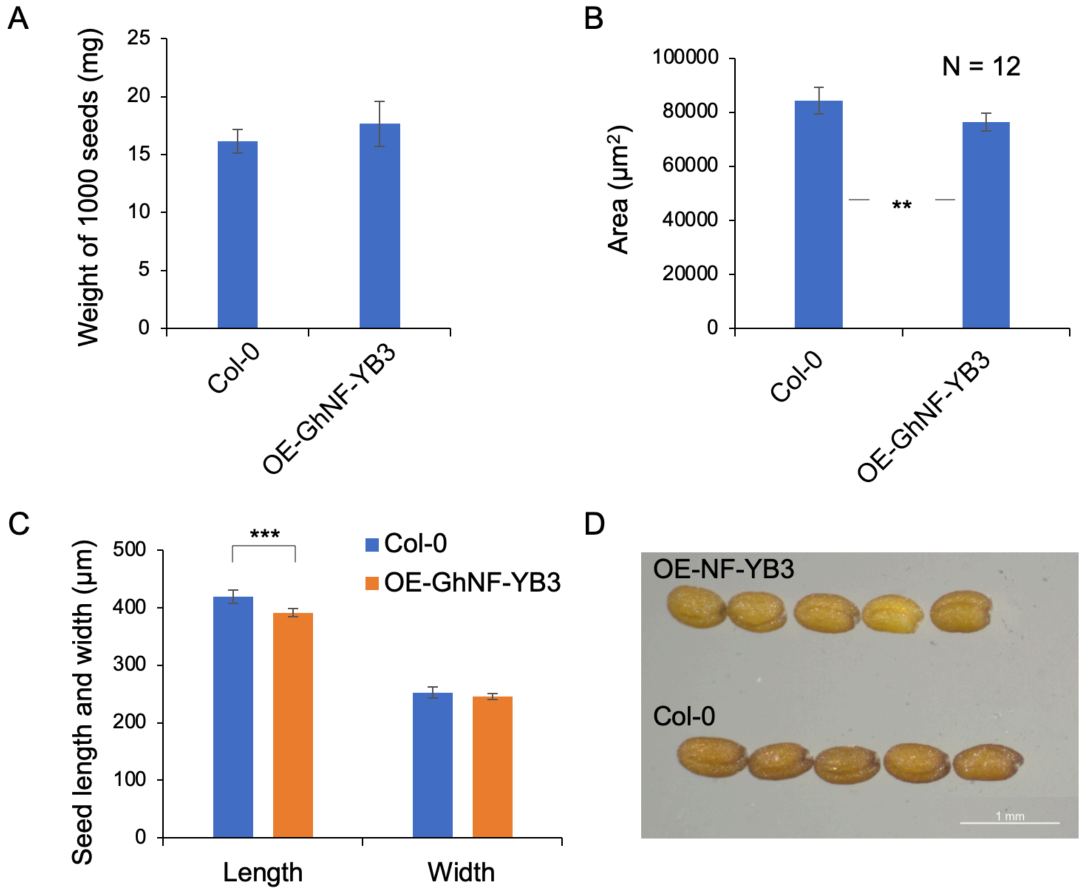


**Figure S13: The phenotypic impacts of the ectopic expression of *GhNF-YB3* in *Arabidopsis*.** The histogram shows the weight of 1000 seeds (A), seed area (B), seed length and width (C) from Col-0 and ectopic expression lines of OE-GhNF-YB3. (D) Photo image shows the seed size difference between the Col-0 and OE-GhNF-YB3. Bar, 1 mm.

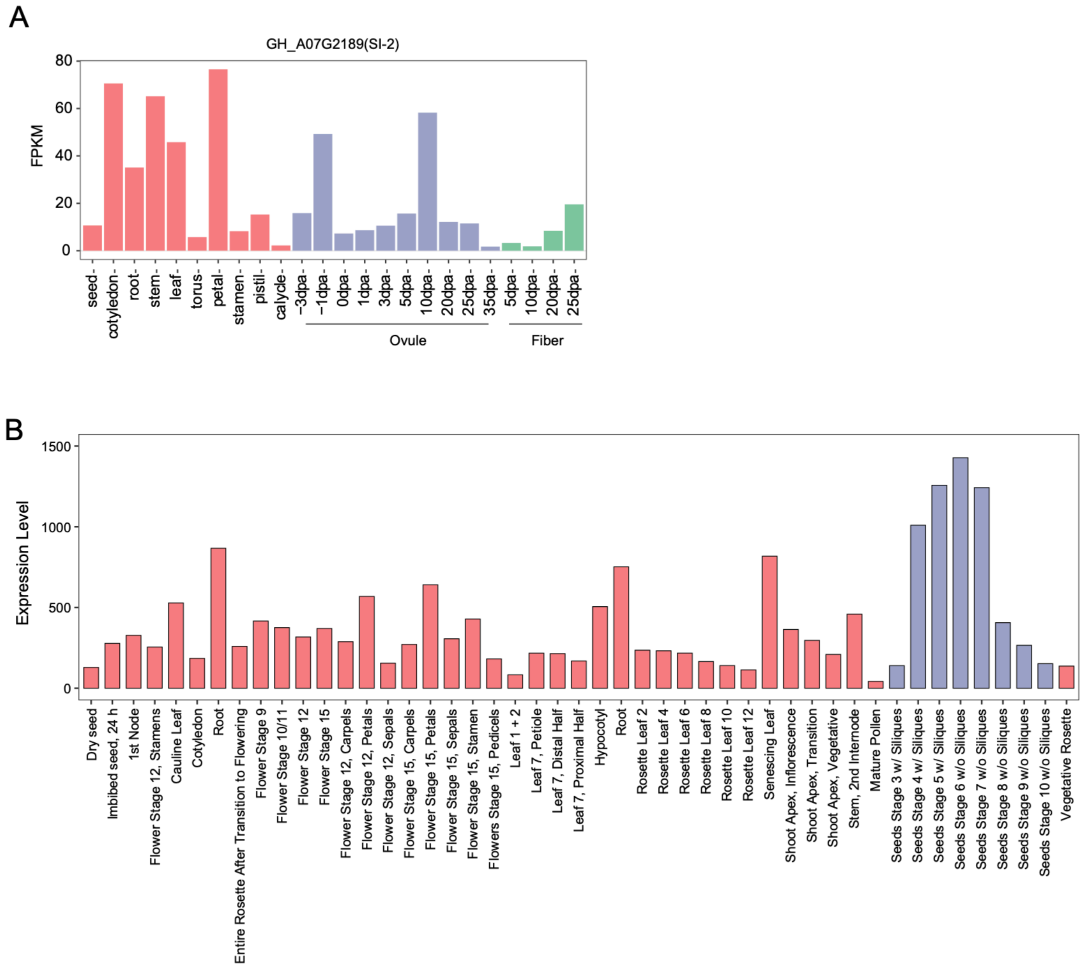


**Figure S14: Expression pattern of *FLA2* homologs in cotton in cotton and *Arabidopsis* tissues.** A: FPKM of *FLA2/GH_A072189* in cotton. B: Expression levels of the *AT4G12730*, which is the orthologous genes of *FLA2/GH_A07G2189* in tissues of *Arabidopsis*. The expression information was obtained from TAIR database (https://www.arabidopsis.org/).
